# Cell-cycle control of chromatin creates a therapeutic window for epigenetic therapy in tumors

**DOI:** 10.64898/2026.09.03.748889

**Authors:** Kourosh Hayatigolkhatmi, Riccardo Valzelli, Elena Ceccacci, Amir Hosseini, Oualid El Menna, Mauro Romanenghi, Isabella Pallavicini, Emanuel Soda, Ornella Ejlli, Emanuela Villa, Paolo Falvo, Pedro Vizan, Enrique Blanco, Sergi Aranda, Roxana Barkhordar, Chiara Soriani, Simona Ronzoni, Giuseppina Giardina, Simona Rodighiero, Francesco Bertolini, Antonello Mai, Sara Polletti, Luisa Lanfrancone, Roberta Noberini, Giuseppe Leuzzi, Pier Giuseppe Pelicci, Tiziana Bonaldi, Luciano Di Croce, Saverio Minucci

## Abstract

The relationship between cell cycle length and differentiation competence has been studied in developmental biology, particularly in embryonic stem cells, which show a very short G1 phase required to maintain pluripotency. How and whether this principle operates in cancer— where the cell cycle is deregulated and G1 is frequently shortened — and whether it can be pharmacologically exploited for therapy, has not been explored.

Here we show that the duration of G1 is a causal determinant of chromatin state in cancer cells and that extending G1 creates a therapeutic window for epigenetic drugs. Using acute myeloid leukemia (AML) as a model, we demonstrate that low-dose palbociclib — at concentrations well below those required for cytostatic arrest — extends G1 without halting proliferation. This modest prolongation reshapes the histone modification landscape: repressive marks (H3K9me2/3, H4K20me2/3) increase while acetylation decreases, and chromatin accessibility rises broadly in the euchromatic compartment. Naturally slow-cycling AML lines share this epigenetic signature regardless of their oncogenic driver mutations, and pharmacologically extending G1 in fast-cycling cells recapitulates it — establishing G1 length as a causal regulator of the cancer epigenome rather than a passive correlate.

To identify epigenetic vulnerabilities created by G1 extension, we performed complementary drug and CRISPR-Cas9 screens in G1-extended AML cells. Both approaches converged on the histone demethylase LSD1 (KDM1A): slow-cycling AML cells are intrinsically sensitive to LSD1 inhibition, while fast-cycling cells become sensitive when G1 is prolonged. The combination of low-dose palbociclib and LSD1 inhibition triggers differentiation and significantly prolongs survival in AML xenograft models.

p21 (CDKN1A) emerges as the central molecular determinant of this response. In slow-cycling AML cells, p21 is highly expressed and its knockdown abolishes LSD1 inhibitor sensitivity. Structure-function analysis using p21 mutants separates the two known activities of p21: the CDK-inhibitory function (which extends G1) is required for sensitization, whereas the PCNA-binding function is dispensable. Three pharmacological routes converge on the same endpoint — CDK inhibition, G1 extension, and a differentiation-competent chromatin state: direct CDK4/6 inhibition by palbociclib, p21 overexpression, and p21 induction through treatment with HDAC or EZH1/2 inhibitors. Palbociclib bypasses the requirement for p21 entirely, confirming that G1 length itself — not p21 as a protein — is the critical variable.

Mechanistically, the combination of G1 extension and LSD1 inhibition yield a qualitatively distinct chromatin state rather than an additive one. ATAC-seq reveals thousands of combination-exclusive accessible regions, enriched for footprints of myeloid differentiation transcription factors including SPI1/PU.1, IRF1, and STAT1/2. A double-lock principle governs this remodeling: palbociclib drives the removal of repressive marks (H3K9me3 and H3K27me3), while LSD1 inhibition installs active marks at the newly accessible regions. The ncBAF chromatin remodeling complex, identified in our CRISPR screen and validated by knockout of its essential subunits BRD9 and SMARCD1, is specifically required for this response. Loss of ncBAF abolishes the combination-induced chromatin remodeling and differentiation program but does not affect the initial G1 extension or retinoic acid–induced differentiation, indicating that ncBAF specifically couples cell-cycle modulation to chromatin remodeling rather than acting as a general differentiation factor.

The principle generalizes beyond AML. In melanoma, breast cancer, and small-cell lung cancer (SCLC), sensitivity to LSD1 inhibition tracks with p21 expression and cycling speed. Primary melanoma samples stratified by p21 expression recapitulate the same pattern: p21-high, slow-cycling cells are sensitive; p21-low, fast-cycling cells are resistant but can be sensitized by palbociclib cotreatment. Cisplatin-induced drug-tolerant persister (DTP) cells — which emerge as a slow-cycling, chemo-resistant population and upregulate both p21 and LSD1 — become vulnerable to LSD1 inhibition and are eradicated by the combination. In melanoma patient-derived xenograft (PDX) models, p21-high tumors respond to LSD1 inhibitor monotherapy, while p21-low tumors are sensitized by palbociclib cotreatment, with p21 knockdown abolishing the response.

Together, these findings establish cell-cycle duration as a tunable regulator of the cancer epigenome and demonstrate that pharmacological G1 extension converts cytostatic CDK4/6 inhibition into an epigenetic sensitization strategy. Both fast-proliferating and slow-cycling tumor compartments — including drug-resistant persister cells — can be targeted by matching the epigenomic state to the appropriate combination of cell-cycle modulators and epigenetic drugs. p21 emerges as a candidate biomarker for patient stratification. More broadly, our work repositions the cell cycle from a passive conduit for proliferation signals to an active, druggable regulator of chromatin fate, with implications that extend from cancer therapy to stem cell biology and regenerative medicine.

## Introduction

The relationship between cell proliferation and differentiation is one of the most fundamental problems in cell biology. Early evidence that the cell cycle is not merely a vehicle for duplication, but an active determinant of cell fate emerged from studies of developmental systems. In the developing rat retina, progenitor cell cycles lengthen as neurogenesis proceeds, and only slowly cycling progenitors give rise to certain neuronal subtypes ^1–3^. These and related observations established the principle that cell cycle length, not merely exit from the cycle, influences the differentiation competence of progenitor cells ^4–6^. However, this concept has remained largely confined to developmental biology, and poorly explored mechanistically.

The strongest mechanistic evidence linking G1-phase length to differentiation competence comes from embryonic stem cells (ESCs). A short G1 phase is an intrinsic determinant of naïve ESC pluripotency, and G1 lengthening is tightly coupled to the exit from pluripotency and the onset of lineage commitment ^7,8^. G1 cyclins directly link proliferation to pluripotency and differentiation programs ^9^. Bivalent epigenetic domains — hallmarks of poised lineage genes — are resolved in a cell-cycle-dependent manner, with G1 lengthening permitting the epigenetic priming that precedes differentiation ^10^. However, ESCs have an atypical cell cycle characterized by truncated G1 and G2 phases, constitutive CDK2 activity, and a weakened G1/S checkpoint. These features are not representative of the cell cycle regulation found in somatic cells, including cancer cells ^11,12^. Whether G1 length controls differentiation competence in cells with a normal or cancer-deregulated cell cycle remains almost entirely unknown.

A limited number of studies have begun to uncover the mechanistic basis for the link between cell cycle machinery and the epigenome, pointing toward direct phosphorylation of epigenetic regulators by cyclin-dependent kinases. CDK1 phosphorylates writers and erasers of all major histone marks, thereby controlling the global epigenetic landscape ^13^. Separately, Polycomb-targeted genes are transcriptionally upregulated during S phase, suggesting that cell cycle phase-specific mechanisms regulate epigenetic state ^14^. These studies provide evidence that the cell cycle machinery communicates with the epigenetic landscape, but they share critical limitations: they were performed exclusively in ESCs, and in most cases are descriptive and did not test whether pharmacologic/genetic modulation of cell cycle is sufficient to reshape the chromatin landscape in a way that licenses differentiation. Additionally, only a few studies address specifically G1 ^15^.

Cancer is fundamentally a disease of cell cycle deregulation. Decades of work have established that tumor cells accumulate mutations driving constitutive proliferation and defective checkpoint responses, and cell cycle kinases are among the most successfully targeted proteins in oncology ^16–19^. Cancer cells frequently exhibit shortened G1 phases, which can promote oncogene-induced DNA replication stress ^20^, and the paths through G1-S are remarkably diverse across tumor types ^21^. Yet the concept that cell cycle length — not merely arrest — controls the differentiation competence and epigenetic state of cancer cells has not been explored. Slow-cycling cancer cells exist in many tumors and are associated with chemoresistance and disease relapse ^22,23^, but their epigenetic and differentiation state has not been systematically characterized. CDK4/6 inhibitors, including Palbociclib, Ribociclib, and Abemaciclib, are FDA-approved drugs that arrest the cell cycle in G1 by inhibiting cyclin D-CDK4/6 complexes ^18,24,25^. Their clinical use is predicated on cytostasis — halting proliferation. However, their mechanism of action is more complex than simple arrest ^26,27^. At sub-maximal doses, CDK4/6 inhibitors could be used not to arrest the cell cycle but to modulate G1 length — extending the time cells spend in G1 without blocking proliferation entirely ^28^. No study has tested CDK4/6 inhibitors as G1-length modulators for epigenetic reprogramming rather than as cytostatic agents ^29,30^.

Acute myeloid leukemia (AML) offers a unique tumor system where differentiation blockade, cell cycle heterogeneity, and epigenetic therapy converge. Leukemic blasts are trapped in an immature state, and the disease is defined by the failure of myeloid differentiation, which is in several cases due to direct action of AML-associated fusion proteins acting as altered transcription factors ^31–34^. AML cell lines span a range of cycling speeds and leukemogenic drivers, providing a tractable system to manipulate cell cycle length. Cell cycle restriction by the endogenous regulator of G1 length p21 matters for self-renewal ^35^, and positive feedback between the transcription factor SPI1/PU.1 and the cell cycle controls myeloid differentiation ^15^, but whether G1 length directly controls the epigenome and differentiation competence of leukemic cells has not been tested. A recent study showing that dual targeting of CDK6 and LSD1 is synergistic in AML ^36^ further supports the convergence of cell cycle and epigenetic mechanisms. Here we set out to systematically investigate whether cell cycle length (focusing on G1) controls the epigenome and differentiation competence of cancer cells, and whether this relationship can be exploited therapeutically.

## Results

### G1-phase length shapes the epigenome of AML cells

Automated single-cell live imaging has shown that low doses of the FDA-approved CDK4/6 inhibitor Palbociclib can extend G1 in tumor cells without inducing cell-cycle arrest. In NB4 AML cells, Palbociclib at concentrations of 50 nM or lower extends G1 without inducing cell-cycle arrest and has minimal effects on the other phases (Fig. 1a; Extended Data Fig. 1a, b) ^28^. Under these conditions, nearly 70% of NB4 cells completed at least one full cell cycle within the first 24 hours, with a significantly extended G1 phase (Extended Data Fig. 1b, c; G1-phase length from 9.1 to 14.2 hours on average; linear mixed-effects model, n = 744 cells from 3 independent experiments, P < 0.0001). Following treatment with 50 nM Palbociclib, NB4 cells display a cell-cycle profile comparable to that of the slow-cycling AML lines Kasumi-1 and UF-1 (Extended Data Fig. 1d) ^37,38^. We therefore established a cellular model in which to ask whether selective G1 prolongation is associated with epigenomic changes in AML cells.

**Figure 1.**
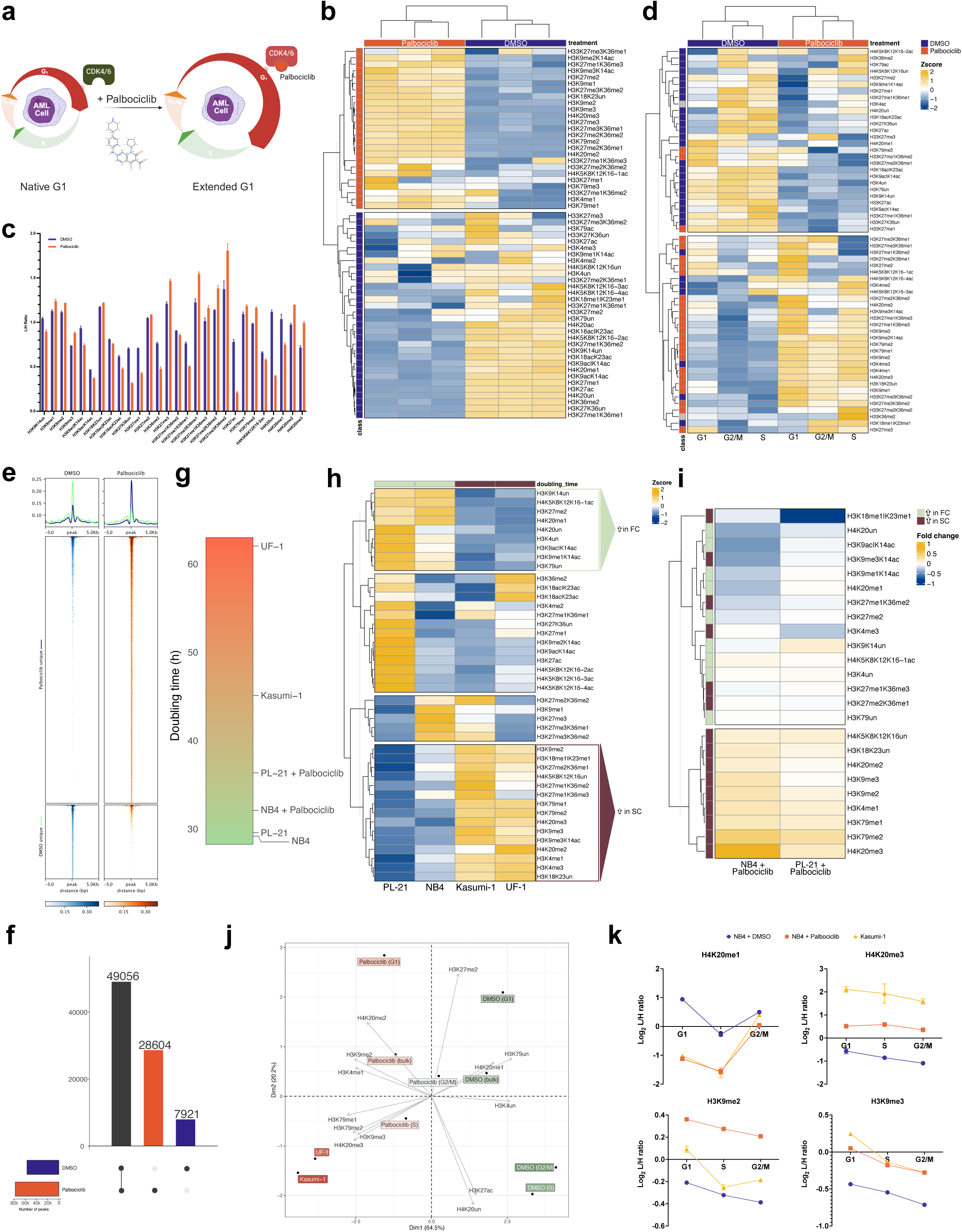
Modulation of cell-cycle length reshapes the epigenetic landscape of AML cells. **a,** Schematic representation of the rationale for using suboptimal doses of Palbociclib to prolong G1 (created with BioRender.com). **b**, Heatmap of MS-quantified hPTMs in NB4 cells treated with DMSO or Palbociclib (50 nM, 24 h). Values are log₂[light:heavy] (L/H) ratios from three biological replicates. Rows (hPTMs) and columns (replicates) were hierarchically clustered; rows further partitioned by k-means (k = 2). Annotation bar highlights hPTMs enriched with Palbociclib (orange) or DMSO (dark blue). Heatmap colors denote log₂[L/H] magnitude (blue→yellow). **c**, Bar chart of significantly altered hPTMs (Statistical significance was assessed using an unpaired two-tailed t-test with Welch’s correction. Multiple comparisons were controlled using the two-stage Benjamini–Krieger–Yekutieli FDR procedure, Q = 1%) in NB4 cells treated as in panel b. Bars represent mean L/H ratios (n = 3); error bars indicate replicate variability. Colors: DMSO (blue), palbociclib (orange). **d**, Heatmap of mean MS log₂[L/H] ratios (n = 3) across G1, S, and G2/M phases in NB4 cells treated with DMSO or Palbociclib (50 nM, 24 h). hPTM class colors as in panel b. **e**, Tornado plots of gained (orange) or lost (dark blue) ATAC-seq signals in G1-sorted NB4 cells treated with Palbociclib (50 nM) versus DMSO for 48 h. Peaks are centered and extended ±5 kb. **f**, UpSet plot of unique and shared ATAC-seq signals from the same experiment. **g**, Heatmap of doubling times (hours) for NB4 and PL-21 cells treated with DMSO or Palbociclib (50 nM and 40 nM, respectively), compared with untreated Kasumi-1 and UF-1 over 6 days. **h**, MS log₂[L/H] ratios (mean of n = 3) in untreated PL-21, NB4, Kasumi-1, and UF-1 cells, hierarchically clustered and partitioned by k-means (k = 4). Side annotation marks clusters enriched in fast-cycling (FC, light green) or slow-cycling (SC, dark red) lines. **i**, FC/SC subsets from “h,” shown for NB4 and PL-21 cells after Palbociclib treatment (50/40 nM, 72 h) as log₂ fold change relative to DMSO (mean of n = 3). **j**, PCA biplot of hPTM profiles (mean log₂[L/H], n = 3). Points represent FUCCI(CA)2-NB4 cells treated with DMSO or Palbociclib (50 nM, 72 h) sorted into G1, S, G2/M as described, or bulk, plus untreated Kasumi-1 and UF-1. Gray arrows indicate hPTM loadings. Dim1 + Dim2 explain ∼90% of variance. Arrow orientation denotes correlation; proximity indicates similar hPTM profiles. **k**, Phase-sorted profiles of four representative hPTMs (H4K20me1, H4K20me3, H3K9me2, H3K9me3). Points show mean log₂[L/H] ratios (n = 2/3) and error bars indicate replicate variability in NB4 cells treated with DMSO (purple, 72 h) or Palbociclib (50 nM, orange, 72 h), and in untreated Kasumi-1 cells (yellow).

We performed unbiased mass spectrometry-based quantification of histone post-translational modifications (hPTMs) in NB4 cells, either untreated or treated with 50 nM Palbociclib for 24 and 72 hours. These analyses revealed that cell-cycle extension induces reproducible changes in hPTM abundance (Fig. 1b, c; Extended Data Fig. 1e, f; Welch’s t-test, Benjamini– Hochberg-adjusted P < 0.01, FDR 1%) ^39–41^. Two distinct clusters of histone modifications emerged: repressive marks, including H3K9me2/3, H3K27me2/3, and H4K20me3, were consistently increased, whereas permissive marks, including H3K27ac, H3K9ac, H3K14ac, and H4K16ac, were reduced. Because Palbociclib-treated NB4 cells contain a higher fraction of cells in G1 than untreated cells (Extended Data Fig. 1d), we sorted NB4 cells stably expressing cell-cycle-stage-specific fluorescent reporters according to cell-cycle phase. Notably, the observed changes were not restricted to G1 but were detected across all cell-cycle phases, demonstrating that they were not caused by cell-cycle-distribution artifacts (Fig. 1d). The sparing of H3K27me3 is consistent with its distinct propagation kinetics: unlike most methylation marks, H3K27me3 is re-established progressively on both new and parental histones over multiple cell generations, making it inherently less sensitive to G1 extension within a single cell cycle ^42,43^.

Analysis of chromatin accessibility using ATAC-seq revealed that hPTM changes are accompanied by global changes in chromatin accessibility (Fig. 1e, f). Chromatin accessibility changes in G1-extended cells were predominantly characterized by de novo or enhanced openings, with relatively few regions showing decreased accessibility (Fig. 1f; Extended Data Fig. 1g). Although the global increase in repressive histone marks might appear at odds with the net gain in chromatin accessibility, the two observations likely reflect changes in distinct genomic compartments. The marks that increase upon G1 extension — H3K9me2/3 and H4K20me2/3 — are predominantly associated with constitutive heterochromatin, which is already poorly or not accessible and therefore invisible to ATAC-seq. In contrast, the accessibility gains observed by ATAC-seq should occur in the euchromatic compartment.

We next analyzed three additional AML cell lines — UF-1 and Kasumi-1 (relatively slow-cycling) and PL-21 (relatively fast-cycling) — which harbor distinct leukemogenic driver mutations ^37,38,44^. Low-dose Palbociclib slowed proliferation of PL-21 cells, mirroring its effect on NB4 cells (Extended Data Fig. 1h). We therefore established six cell-cycle models: two naturally slow-cycling lines, two naturally fast-cycling lines, and the two fast-cycling lines under G1-prolonging low-dose Palbociclib (Fig. 1g). Mass spectrometry profiling of hPTMs showed that UF-1 and Kasumi-1 share an epigenetic signature distinct from that of NB4 and PL-21 (Fig. 1h) — despite carrying different oncogenic drivers — indicating that this signature tracks with cycling speed rather than genotype. Fast-cycling lines were enriched for acetylated and unmodified histone residues, consistent with rapid histone turnover, whereas slow-cycling lines were enriched for methylation states including H3K9me2/3, H4K20me2/3, H3K79me1/2, H3K4me1/3, and H3K27me3 — marks linked to heterochromatin assembly, DNA-damage signaling, replication timing, and regulatory-element activity ^45–52^. Notably, extending G1 in fast-cycling cells increased hPTMs characteristics of slow-cycling cells and decreased those of fast-cycling cells (Fig. 1i): histone acetylation fell, while H3K9me2/3, H4K20me2/3, and H3K79me1/2 rose. The notable exception was H3K27me3: although enriched in naturally slow-cycling lines, this Polycomb-linked mark was not increased by G1 extension. Thus, G1 prolongation does not uniformly raise repressive marks but selectively engages the H3K9 and H4K20 methylation pathways while leaving Polycomb-mediated repression unchanged — pointing to directed rather than passive epigenetic remodeling.

To exclude confounding by cell-cycle-dependent oscillation of hPTMs, we sorted cells into G1, S, and G2/M fractions and reanalyzed hPTMs by mass spectrometry ^42,53^. Principal component analysis (PCA) of the most consistently altered hPTMs revealed that Palbociclib-treated NB4 cells shift along the first component (64.5% of variance) toward the chromatin profile of the naturally slow-cycling lines Kasumi-1 and UF-1, irrespective of cell-cycle phase (Fig. 1j). This shift was driven by increased methylation and reduced acetylation marks, confirming the previous analysis. Quantification of L/H ratios for representative marks showed that Palbociclib-treated NB4 cells approach or match the Kasumi-1 pattern in every phase (Fig. 1k), indicating a stable epigenetic change in decelerated cells rather than a G1-restricted artifact or complete arrest. Crucially, because pharmacologic G1 extension in fast-cycling cells recapitulates the core features of the natural slow-cycling signature, G1 length is causal in shaping the epigenome rather than being merely associated with it. Together, these findings establish that G1 prolongation reshapes the epigenomic landscape of AML cells — hPTMs and chromatin accessibility — directly linking cell-cycle length to epigenetic state.

### Cell cycle prolongation determines response to epigenetic therapy in AML

Given the observed epigenetic changes upon cell-cycle prolongation, we explored whether low doses of Palbociclib treatment would create dependencies on epigenetic factors, thereby revealing potential combination strategies with epigenetic drugs (epi-drugs). We followed two complementary strategies in fast-cycling AML NB4 cells treated with low-dose Palbociclib: a focused drug screen of 290 epigenetic modulators and a CRISPR-Cas9 screen targeting 652 chromatin-related genes (Fig. 2a). Hits were prioritized as targets that demonstrated cooperation between small-molecule and sgRNA targeting upon G1 prolongation. Across both approaches, G1-extended AML cells showed a marked response to LSD1 (*KDM1A*) perturbation in G1-extended AML cells (Fig. 2b, c). LSD1, mainly responsible for demethylating activities on H3K4/K9me1/2, is a transcriptional (co-)regulator that is often overexpressed in AML ^54–56^. Previous studies from our laboratory and others have extensively characterized how deregulated catalytic and scaffolding functions of LSD1 contribute to leukemogenesis by supporting differentiation blockade ^54,57,58^. Therapeutic strategies targeting LSD1, both as single agents and in combination with other treatments, have been evaluated across various leukemic models to overcome this block ^54,57–61^, and LSD1 functions as a co-regulator of key hematopoietic transcription factors (TFs) such as GFI1(B) and SPI1 ^58,60^. We therefore chose LSD1 inhibition as a model to characterize the mechanism of enhanced response to epigenetic therapy in cell cycle–modulated AML cells.

**Figure 2.**
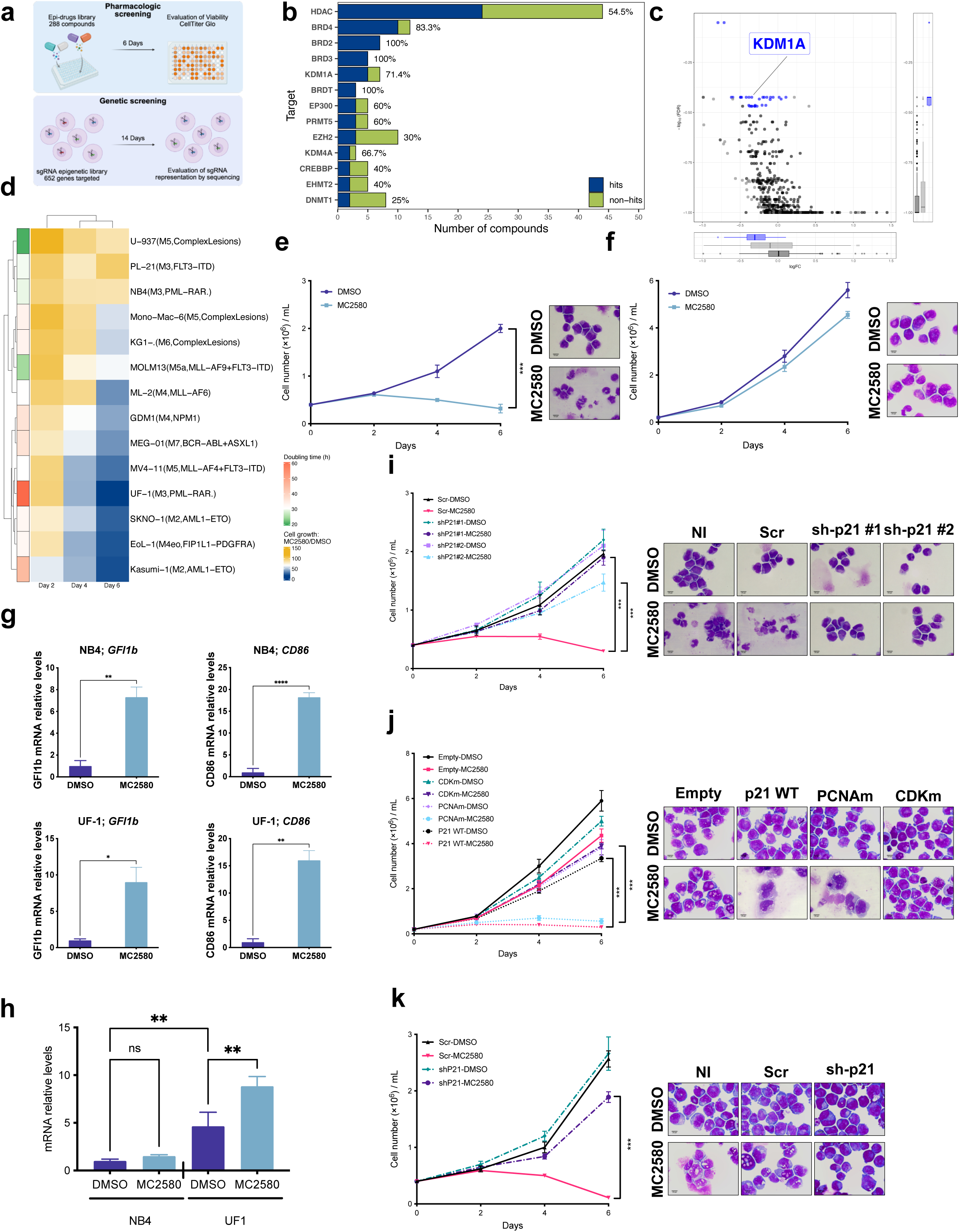
AML cell response to LSD1 inhibition is determined by G1 length. **a,** Schematic representation of pharmacologic (top) and genetic CRISPR drop-out (bottom) screening approaches (created with BioRender.com). **b**, Bar chart of epigenetic targets identified from a 290-compound library. Hits were defined as compounds reducing NB4 cell viability to ≤30% of the 50 nM palbociclib control after 6 days of treatment with 230 nM of each compound (average estimated IC70) plus 50 nM palbociclib (CTG readout). Bars indicate the total number of compounds tested for each target, partitioned into hits (blue) and non-hits (green), with percentages indicating the fraction of hits. For visualization, the plot was restricted to targets represented by at least two hit compounds. **c**, Scatter plot ranks the target genes based on the “negative” (DOWN) log_2_FC of all the 6 sgRNAs and the 3 replicates, comparing 14 days DMSO- or 50 nM palbociclib-treated cells. The top 30 genes targeted by sgRNAs that are under-represented in palbociclib-treated samples are highlighted in blue. **d**, Heatmap of cell counts in a panel of FAB-comprehensive AML cell lines treated with MC2580 (2 µM, 6 days), normalized to DMSO. Hourly doubling times under normal growth conditions are color coded. **e–f**, Growth curves (6 days) and May-Grünwald-Giemsa-stained images (day 7) of NB4 (e) and UF-1 (f) cells treated with DMSO or MC2580 (2 µM). Data are mean ± SD of triplicates. **g**, Bar charts showing analysis of *GFI1b* and *CD86* mRNA relative levels in NB4 and UF-1 cells treated with DMSO or MC2580 (2 µM). Values were normalized to *GAPDH* relative to DMSO-treated. Data are presented as mean ± SD. P values were obtained using a two-tailed unpaired t-test (ns, not significant; ✱P < 0.05). **h**, Bar chart showing analysis of *CDKN1A* mRNA relative levels in NB4 and UF-1 cells treated with DMSO or MC2580 (2 µM). Values were normalized against *GAPDH*. Data are presented as mean ± SD. Statistical significance was assessed by ordinary one-way ANOVA followed by Tukey’s multiple-comparisons test. ns, not significant; ✱P < 0.05, ✱✱P < 0.01, ✱✱✱P < 0.001, ✱✱✱✱P < 0.0001. **i–k**, Growth curves (6 days) and May-Grünwald-Giemsa images (day 7) of UF-1 (i), NB4 (j), and Kasumi-1 (k) cells stably transduced with the indicated vectors and treated with DMSO or MC2580 (2 µM). Data are mean ± SD of triplicate. All quantitative growth data across the figure were analyzed by one-way ANOVA with Bonferroni’s post-hoc test. P < 0.05 (✱), P < 0.01 (✱ ✱) and P < 0.001 (✱ ✱ ✱).

We next treated a comprehensive panel of AML cell lines — spanning FAB classes M2 through M7 (6 of 8 FAB classes) — with the irreversible Tranylcypromine (TCP)-based LSD1 inhibitor MC2580 ^55,62^. We observed a spectrum of responses, with the fastest-cycling models showing the highest levels of resistance to LSD1 inhibition (Fig. 2d; Extended Data Fig. 2a). UF-1 cells, which harbor the same leukemogenic driver mutation as NB4 cells, were the slowest cycling model in this panel and showed the highest sensitivity to LSD1 inhibition, leading to impaired proliferation and morphological differentiation, unlike NB4 cells (Fig. 2e, f; Extended Data Fig. 2b). Two additional LSD1 inhibitors, GSK-LSD1 and DDP38003, similarly suppressed UF-1 proliferation (Extended Data Fig. 2c, d), and shRNA-mediated knockdown (KD) of LSD1 in UF-1 cells reduced cell proliferation and induced differentiation (Extended Data Fig. 2e–h), confirming the on-target mechanism of action. A colony-forming unit (CFU) assay showed that MC2580 strongly reduced UF-1 colony formation, yielding small, diffuse colonies instead of the large, compact colonies seen in controls (Extended Data Fig. 2i, j). Immunoblotting showed similar LSD1 levels in NB4 and UF-1 cells, ruling out basal expression as a determinant of sensitivity (Extended Data Fig. 2k). Moreover, MC2580 increased H3K4me2 and upregulated LSD1-repressed genes (*GFI1B* and *CD86*) in both UF-1 and NB4 cells, indicating effective LSD1 inhibition in both models (Fig. 2g; Extended Data Fig. 2l) ^63,64^.

RNA-seq revealed a high transcriptional correlation between UF-1 and NB4 cells, yet pathway analysis on the few differentially expressed genes (DEGs) showed distinct programs driven by TFs such as SPI1 and MYC and differential regulation of G1-to-S transition genes (Extended Data Fig. 2b, m) ^31,38^. Among these, *CDKN1A* (encoding p21), a negative regulator of G1-to-S progression ^65^, was significantly upregulated in the slow-cycling UF-1 cells (Extended Data Fig. 2n; [logFC = 5.15; FDR = 6.4 × 10⁻⁵]).a MC2580 treatment further increased p21 expression and the proportion of cells in G1 in UF-1 cells, but not in NB4 cells (Fig. 2h; Extended Data Fig. 3a, b). We used a genetic approach to knock down p21 in UF-1 cells using two distinct shRNA constructs (Extended Data Fig. 3c). In p21-KD UF-1 cells, we observed a reduction in the G1 proportion and an increase in the S-phase fraction (Extended Data Fig. 3d), and the p21 induction upon LSD1 inhibition was abolished (Extended Data Fig. 3e, f). p21 KD countered the growth-inhibitory and differentiation-inducing effects of MC2580 in UF-1 cells (Fig. 2i). These data establish that p21 is necessary and sufficient for LSD1 inhibitor sensitivity within the p21-induction axis in UF-1 cells. Beyond its role in cell cycle regulation, p21 also interacts with PCNA and participates in DNA replication ^65^. To dissect which function of p21 mediates the sensitization, we moderately overexpressed p21 in NB4 cells using three constructs encoding either wild-type (WT) p21 or mutant forms lacking interaction with CDKs (CDKm) or PCNA (PCNAm) (Extended Data Fig. 3g). WT and PCNAm p21 slowed NB4 growth without inducing morphological changes, mimicking the mild effect of Palbociclib, and these cells became sensitized to LSD1 inhibition. In contrast, CDKm-p21-expressing cells did not (Fig. 2j; Extended Data Fig. 3h, i). These results demonstrate that the sensitizing effect of p21 is mediated through its CDK-inhibitory (G1-extending) function, not its PCNA (replication) function, directly linking the effect to the cell cycle regulation.

To extend this observation to other AML subtypes, we knocked down p21 in the slow-cycling Kasumi-1 cell line (Extended Data Fig. 3j), which showed high sensitivity to LSD1 inhibition (Fig. 2d). KD of p21 in Kasumi-1 cells abrogated both growth suppression and differentiation, and blocked p21 upregulation following MC2580 treatment (Fig. 2k; Extended Data Fig. 3k, l), similar to the pattern observed in UF-1 cells.

To pharmacologically mimic p21 induction, we employed HDAC and EZH1/2 inhibitors — previously reported to regulate *CDKN1A* expression ^66–68^. NB4 cells were treated with two HDAC inhibitors, the FDA-approved Vorinostat (SAHA) and Trichostatin A (TSA), for 24 hours, followed by MC2580 (Fig. 3a) ^69^. Concentrations and timing were chosen to achieve a level of p21 upregulation comparable to the basal expression seen in UF-1 cells (Extended Data Fig. 4a). While SAHA and TSA alone had minimal effects on proliferation at the doses tested, they strongly sensitized NB4 cells to MC2580, significantly reducing cell growth and inducing differentiation (Fig. 3b, c). To confirm that this cooperative effect occurs via p21 induction, we knocked down p21 in NB4 cells (Extended Data Fig. 4a, b). p21 knockdown prevented G1 extension, blocked differentiation, and reversed the anti-proliferative effects of HDAC inhibitor and MC2580 cotreatment, establishing p21 as the critical mediator of this combinatorial effect (Fig. 3b, c; Extended Data Fig. 4c).

**Figure 3.**
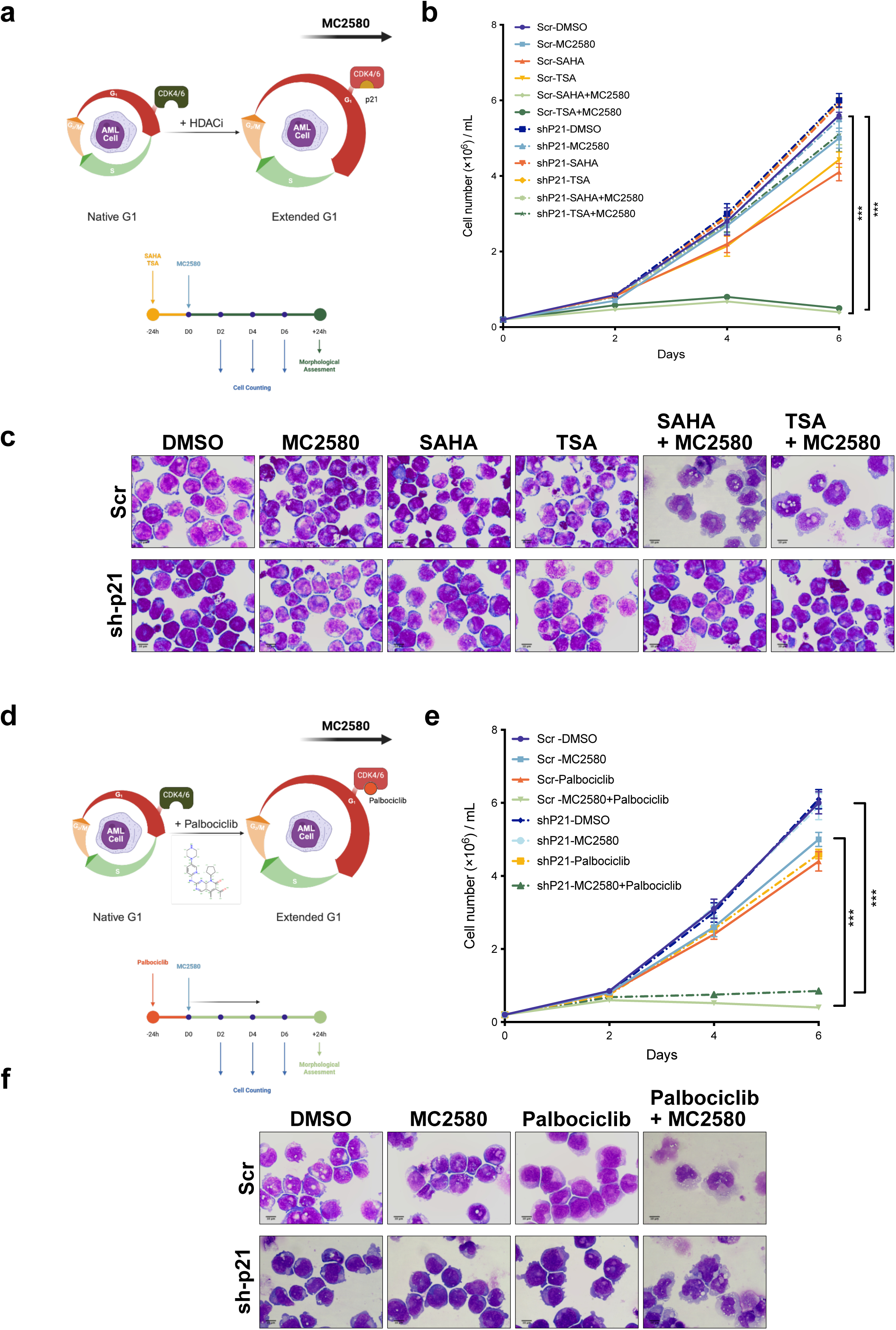
Pharmacological G1 extension sensitizes resistant AML cells to LSD1 inhibition. **a**, Experimental scheme for combining HDAC inhibitors with LSD1 inhibition (created with BioRender.com). **b–c**, Growth curves (6 days) (b) and May-Grünwald-Giemsa images (day 7) (c) of NB4 cells transduced with the indicated vectors and treated with DMSO, MC2580 (2 µM), HDAC inhibitors (SAHA or TSA, 20 nM), or combinations. **d**, Experimental scheme for combining CDK4/6 inhibition (Palbociclib) with LSD1 inhibition (created with BioRender.com). **e–f**, Growth curves (6 days) (e) and May-Grünwald-Giemsa images (day 7) (f) of NB4 cells treated with DMSO, MC2580 (2 µM), Palbociclib (50 nM), or their combination. Data in b and e are mean ± SD of triplicates. All quantitative growth data across the figure were analyzed by one-way ANOVA with Bonferroni’s post-hoc test. P < 0.05 (✱), P < 0.01 (✱ ✱) and P < 0.001 (✱ ✱ ✱).

We next tested whether removal of H3K27me3 at *CDKN1A* regulatory elements could similarly upregulate p21. Treatment with the EZH1/2 inhibitor UNC1999 — targeting the main H3K27me3 methyltransferases ^70^ — required prolonged exposure (4 days) to erase H3K27me3 and induce p21 (Extended Data Fig. 4d, e). Consistently, UNC1999-mediated p21 induction sensitized NB4 cells to LSD1 inhibition, and this effect was again abolished by p21 knockdown (Extended Data Fig. 4f), confirming that p21 upregulation underlies the response. These data show that pharmacologic induction of p21 can be achieved using different epi-drugs, poising AML cells toward differentiation induction via secondary targeted epigenetic therapies such as LSD1 inhibitors. Together, these findings demonstrate that cell cycle length shapes the response of AML cells to epigenetic therapies, and that p21 is one sufficient, CDK-dependent route to the G1-extended state that licenses this response.

We then tested whether G1-phase lengthening through pharmacologic CDK4/6 inhibition could recapitulate the synergistic effects observed with p21-driven G1 extension. As previously shown, 24-hour treatment with 50 nM Palbociclib in NB4 cells shifted their cell-cycle profile to resemble that of slower-cycling Kasumi-1 and UF-1 cells (Extended Data Fig. 1c, d) ^69,71^, accompanied by modulation of hPTMs toward the slow-cycling-specific epigenetic state (Fig. 1h, i). We designed a treatment schedule where NB4 cells were first exposed to 50 nM Palbociclib for 24 hours, followed by MC2580, in cells expressing either control or p21-targeting shRNAs (Fig. 3d). Palbociclib sensitized NB4 cells to LSD1 inhibition — modulating the cell cycle status, reducing proliferation, and inducing differentiation — regardless of p21 expression (Fig. 3e, f; Extended Data Fig. 4g). This result is mechanistically consistent: since Palbociclib directly inhibits CDK4/6 — the same endpoint achieved by p21 through its CDK-inhibitory function — it bypasses the requirement for p21 induction. The p21 independence of Palbociclib therefore strengthens the conclusion that G1 length, not p21 *per se*, is the critical determinant of LSD1 inhibitor response. Moreover, low doses of other CDK inhibitors, such as the pan-CDK inhibitor Milciclib, similarly sensitized NB4 cells to LSD1 inhibition (Extended Data Fig. 4h), and cell-cycle prolongation by low-dose Palbociclib also sensitized PL-21 cells (driven by a different leukemogenic driver mutation) to LSD1 inhibition (Extended Data Fig. 4i).

To assess *in vivo* efficacy, we established NB4 CDX models in NSG-SGM3 mice. We detected luciferase-labeled NB4 cells in the liver within days of injection and subsequent colonization of the bone marrow within one week (Extended Data Fig. 4j). Such rapid and aggressive AML infiltration resulted in a narrow therapeutic intervention window. Despite this limitation and the reported toxicity of higher doses of DDP38003 (the *in vivo*–suitable analog of MC2580 ^72^) in NSG mice, the combination of Palbociclib and DDP38003 significantly prolonged survival in AML-engrafted mice (Extended Data Fig. 4k; at least 8 mice per group; overall: Mantel–Cox log-rank test, χ²(3) = 16.23, P = 0.0010; combination VS NT: adjusted P = 0.0275; Holm–Šídák correction) ^73^. These findings demonstrate that targeting the cell cycle can prime AML cells for epigenetic therapies, offering customizable combination strategies for diverse genetic contexts.

### G1 extension reprograms AML chromatin toward a differentiation-competent state

The results above establish a unified causal model: G1 length is the upstream determinant of LSD1 inhibitor response. p21 is one sufficient, CDK-dependent route to G1 extension, its sensitizing effect requires its CDK-inhibitory function and p21 is dispensable when CDKs are inhibited directly by Palbociclib. HDACi and EZH1/2i represent a third route, acting through p21 induction to achieve the same CDK inhibition and G1 extension. All three routes converge on the same endpoint: CDK inhibition, G1 extension, and a differentiation-competent chromatin state. This raised a mechanistic question: how does G1 extension reshape the chromatin landscape to enable LSD1 inhibitor–driven differentiation?

G1 extension converts LSD1 inhibition into a differentiation program. Our ATAC-seq data revealed that Palbociclib treatment alone in NB4 cells induces 28604 unique peaks, among which 14,951 (regions MPC and PC) persist upon Palbociclib + MC2580 cotreatment (Fig. 4a). LSD1 inhibition by MC2580 led to 21872 peaks, among which 11,636 (regions MPC and MC) were maintained upon combination treatment. The combination treatment led to 13,853 exclusive open chromatin peaks (region C) (Fig. 4a; consensus peaks detected in ≥3 of 5 biological replicates; MACS FDR < 0.001). Notably, these combination-exclusive regions constitute 43% of all combination-opened chromatin and form the largest single class, while 18,856 regions opened by either single agent (regions P and M) are not reproduced in the combination; the combination therefore establishes a qualitatively distinct chromatin state rather than an additive one. The combination-opened regions are expected to harbor key regulatory loci required for epigenetic and transcriptional reprogramming toward differentiation. ATAC-seq footprinting analysis ^74^ revealed significantly enhanced footprints of several TFs, most prominently the ETS family (SPI1/PU.1), followed by IRF1, STAT1/2, PRDM1, and GFI1(B) upon combination treatment (Fig. 4b; Extended Data Fig. 5a). The region-resolved footprinting pattern mirrors the pharmacological logic of the combination: GFI1/GFI1B footprints mark mainly all MC2580-responsive classes (M, MC, MPC and C) but are less discovered in Palbociclib-only regions, consistent with non-catalytic displacement of LSD1 from GFI1/GFI1B-tethered repressor complexes, whereas SPI1, CEBPA, IRF-family and STAT1/STAT2 footprints dominate the combination-opened classes, including the combination-exclusive C regions ^58,75^. Most of these TFs are well-known regulators of early myeloid differentiation, for example SPI1 and IRF1, which induce granulocytic differentiation and proliferation of hematopoietic cells ^60,76–81^. We next examined whether the increased footprints reflected elevated expression, enhanced chromatin-binding capacity, or both. Most of these TFs showed increased RNA expression, with corresponding protein levels rising concurrently or with a slight delay (Fig. 4c; Extended Data Fig. 5b, c). Strikingly, the transcriptional programs induced by Palbociclib and by the combination were largely non-overlapping: Palbociclib alone induced a progenitor/lineage-priming repertoire (RUNX1, IKZF1, BCL11B, STAT1), whereas only the combination induced the myeloid execution module (SPI1, CEBPA, GFI1/GFI1B, IRF1/IRF2, STAT2, PRDM1, BATF) — the same factors whose footprints mark the combination-opened regions, establishing a self-reinforcing loop between chromatin opening and factor induction ^58,60,77,79,81–83^. In the naturally slow-cycling Kasumi-1 and UF-1 cells, ETS family binding motifs — and SPI1 in particular — were prominently found in mutually open chromatin regions between the native state of these models and G1-extended NB4 cells (Extended Data Fig. 5d). The same motifs were mutually enhanced upon LSD1 inhibition in all three models, suggesting that slowing the cell cycle in fast-cycling cells is a transcriptional regulatory prerequisite for differentiation induction relying on these TFs (Extended Data Fig. 5e). Together, these findings indicate that effective and coordinated recruitment of these factors to chromatin is facilitated by cell-cycle–dependent chromatin opening.

**Figure 4.**
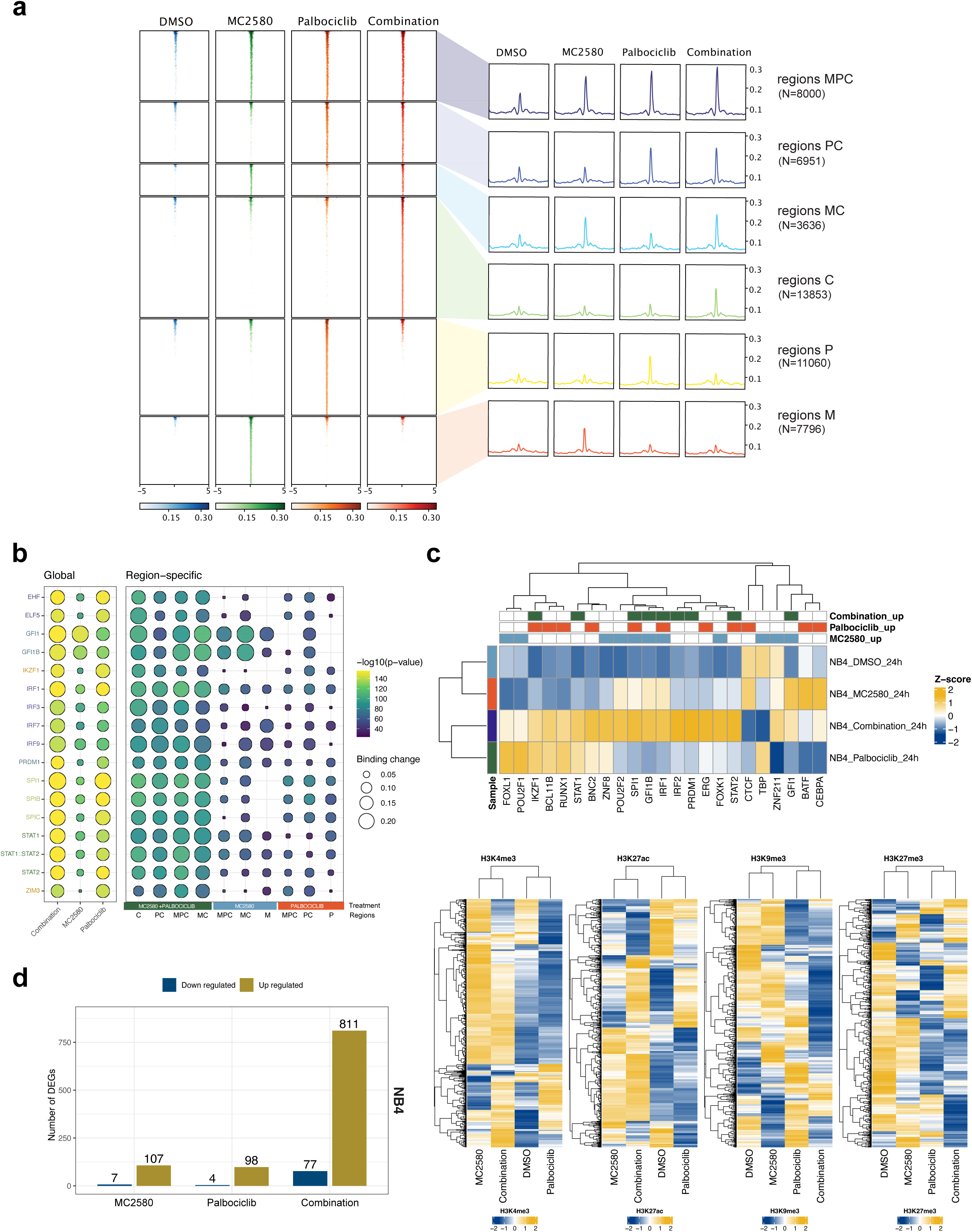
G1 extension primes AML chromatin for differentiation. **a,** Tornado plots (left) of unique and shared ATAC-seq peaks in NB4 cells treated with Palbociclib (50 nM), MC2580 (2 µM), or their combination versus DMSO (day 1), and peak profiles (right) annotated with initials of the indicated treatments. Peak numbers are indicated. **b**, Bubble plot showing TOBIAS footprinting comparing TF binding in NB4 cells treated with Palbociclib + MC2580, Palbociclib, or MC2580 versus DMSO (day 1). Top TFs in combination-related comparisons were selected using the most significant motif per TF. Regions referring to panel “a” are indicated by the same initials. Circle size indicates differential binding and color intensity reflects −log10(P-value); negative differential binding values are not shown.. **c**, Heatmap of RNA expression for the top 20 significantly enriched TFs (positively bound in at least one condition). Top annotation indicates treatment-specific detection. **d**, Bar charts showing DEGs (FDR < 0.01, |FC| > 1.5) in NB4 treated with 2 µM MC2580, 50 nM Palbociclib, or combination (day 1). e, Heatmaps showing the enrichment of four hPTMs (left to right: H3K4me3, H3K27ac, H3K9me3 and H3K27me3) measured by ChIP-seq within −2.5/+2.5 kb of the TSS of 811 genes upregulated upon palbociclib + MC2580 treatment, shown in panel “d.”

We then asked how the combined transcriptional and epigenetic effects of G1 prolongation and LSD1 inhibition leads to differentiation. Hence, we assessed early transcriptional changes in NB4 cells upon treatment with LSD1 inhibitor, Palbociclib, and the combination. The combination treatment induced more DEGs than the two single treatments combined, suggesting an enhanced transcriptional response (Fig. 4d). Comparing these changes with those in genetically distinct PL-21 cells under the same treatment schedule revealed highly similar early transcriptional responses, reducing the transcriptional discrimination between the two models (Extended Data Fig. 5f). Pathway analyses showed that these genes are involved in inducing similar differentiation pathways, particularly activating granulocytic differentiation cascades and immune response (Extended Data Fig. 5g). To investigate the epigenetic state associated with the upregulated genes, we performed ChIP-seq for four key hPTMs at their loci: H3K4me3 (active promoters), H3K27ac (active regulatory/enhancer regions), H3K27me3 (facultative heterochromatin), and H3K9me3 (constitutive heterochromatin) (Fig. 4e). Activating marks were increased and repressive marks were decreased upon both treatments. However, LSD1 inhibition was the main driver of the increase in activating hPTMs, while Palbociclib initiated — and the combination completed — the removal of repressive hPTMs. Although bulk mass spectrometry revealed a global increase in repressive marks upon G1 extension (Fig. 1b–d), the locus-specific analysis by ChIP-seq at upregulated genes showed the opposite pattern — loss of repressive marks. As speculated formerly, this apparent divergence reflects the compartment-specific nature of the two measurements: the global increase captured by mass spectrometry is dominated by constitutive heterochromatin, which constitutes the bulk of the genome, whereas the loss observed by ChIP-seq occurs at the regulatory elements of the specific genes targeted by the treatments. The locus-specific loss of H3K27me3 at these differentiation genes, contrasted with the global H3K27me3 sparing observed upon G1 extension (Fig. 1i), further indicates that Polycomb-mediated repression is redistributed away from differentiation loci rather than globally inactivated. This division of labor — Palbociclib initiating chromatin opening by removing repressive marks, LSD1 inhibition depositing activating marks, and the combination completing both — supports the efficacy of the combination regimen in inducing differentiation through a defined epigenetic and transcriptional reprogramming toward the myeloid lineage.

### The G1-extended differentiation-competent chromatin state relies on the ncBAF chromatin remodeler complex

Extending G1-phase length alters global and local hPTM distribution and chromatin accessibility, and, when combined with LSD1 inhibition, amplifies these effects to enhance TF activity and differentiation through altered gene expression. This transition from epigenetic remodeling to transcriptional output likely involves chromatin remodelers and other enzymatic complexes that interpret the histone code to coordinate chromatin reorganization and transcriptional activation. Among the chromatin regulators in our CRISPR screen, sgRNAs targeting three independent subunits of the SWI/SNF complex non canonical BAF (ncBAF) — BRD9, SMARCD1, and ACTL6A — were significantly enriched upon Palbociclib treatment (Fig. 5a), suggesting that ATP-dependent chromatin remodeling complexes contribute to the cellular response to CDK4/6 inhibition. To determine whether ncBAF is actively redistributed on chromatin, we performed BRD9 ChIP-seq, using cells sorted in the G1 phase to exclude differences arising from treatment-induced cell-cycle redistribution. We focused on BRD9 since it is uniquely associated with the ncBAF complex, whereas SMARCD1 and ACTL6A are shared with the cBAF and PBAF complexes. Upon G1-extension, BRD9 occupancy was markedly increased at regulatory regions surrounding the transcription start sites of the combination-upregulated genes identified formerly (Fig. 5b), consistent with active recruitment of the ncBAF complex to remodel chromatin at target gene loci.

**Figure 5.**
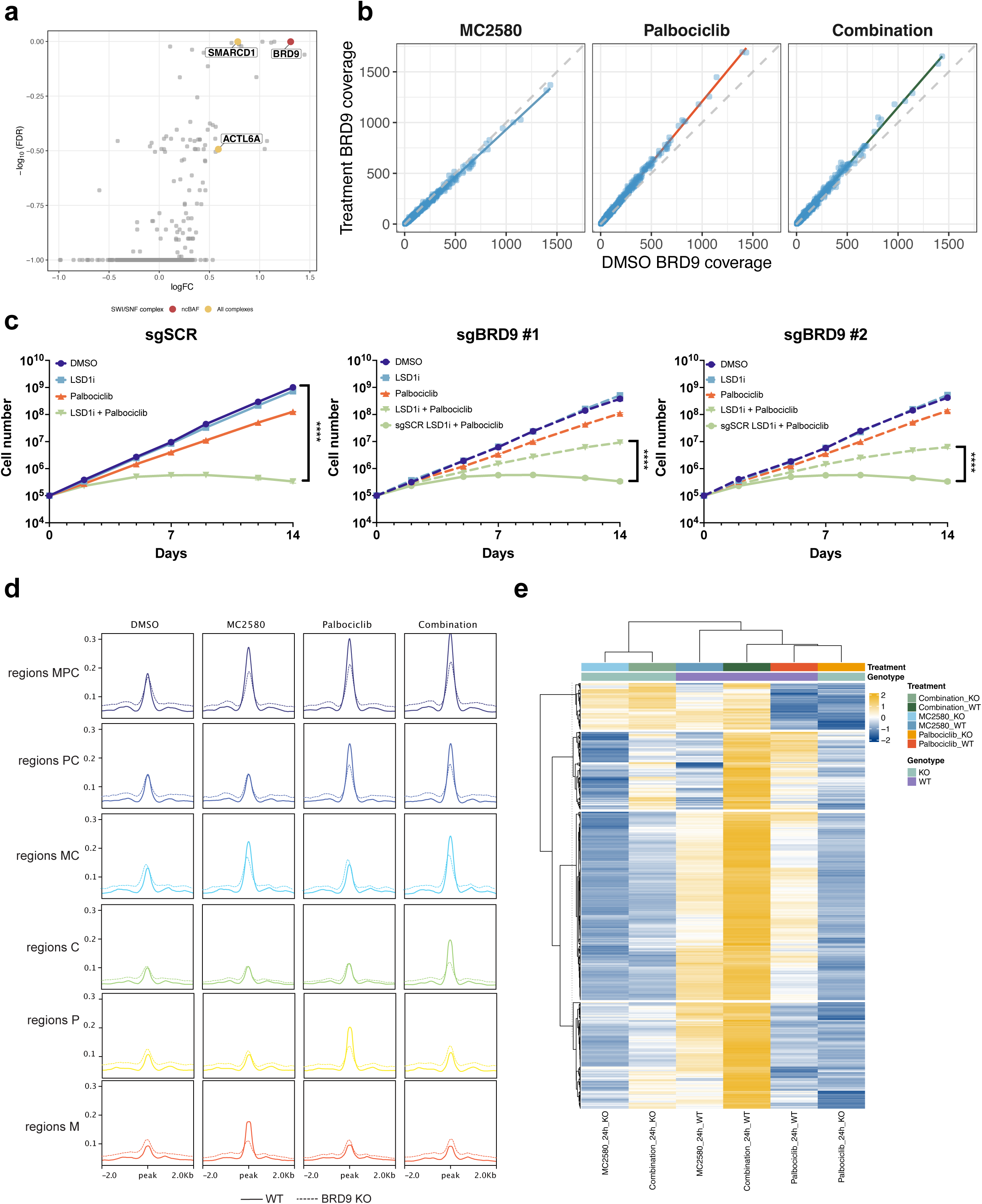
The ncBAF chromatin reader complex mediates chromatin accessibility and differentiation-associated transcription in G1-extended NB4 cells. **a**, Scatter plot showing positive (UP) log_2_FC values from six sgRNAs across three replicates, comparing cells treated with 50 nM palbociclib for 14 days with DMSO-treated cells. BRD9, SMARCD1, and ACTL6A are highlighted according to their SWI/SNF complex membership. **b**, Scatter plots showing BRD9 ChIP-seq enrichment vs DMSO across the loci of the 811 genes upregulated upon combined Palbociclib and MC2580 treatment. **c**, Growth curves (14 days) of sgRNA-expressing NB4 cells treated with DMSO, MC2580 (2 µM), Palbociclib (50 nM), or combination shown in Log10 scale. Data are mean ± SD of triplicates. **d**, Peak profiles of unique and shared ATAC-seq peaks in WT (normal line) and BRD9-KO (dotted line) NB4 cells treated with palbociclib (50 nM), MC2580 (2 µM), or their combination versus DMSO (day 1) and sorted in G1 phase, referring to the same regions in Figure 4a. **e**, Heatmap of RNA expression for the top 811 significantly upregulated genes shown in Figure 4d in WT (normal line) and BRD9-KO (dotted line) NB4 cells treated with palbociclib (50 nM), MC2580 (2 µM), or their combination versus DMSO (day 1). Data are mean ± SD of triplicates. All quantitative growth data across the figure were analyzed by two-way ANOVA with Tukey’s multiple comparisons test. P < 0.05 (✱), P < 0.01 (✱ ✱) and P < 0.001 (✱ ✱ ✱). Analysis of the data in panel c was performed on log_10_-transformed values.

To establish whether ncBAF activity is required for this response, we generated pooled knockout (KO) cell lines targeting either BRD9 or SMARCD1, two essential components of the complex identified in the screen. Efficient loss of both proteins was confirmed by western blotting (Extended Data Fig. 6a). Loss of either subunit resulted in the failure of cells to undergo the stable proliferative arrest observed in control cells under combination treatment; knockout cells continued to expand throughout the two-week treatment period (Fig. 5c; Extended Data Fig. 6b). Notably, this phenotype was not accompanied by detectable alterations in cell-cycle distribution, as DAPI staining revealed comparable profiles between knockout and wild-type cells with or without Palbociclib treatment (Extended Data Fig. 6c), indicating that ncBAF is dispensable for the initial G1 lengthening elicited by CDK4/6 inhibition, but it is necessary for the establishment of the durable growth arrest associated with the differentiation response.

Although proliferation was partially restored in BRD9- and SMARCD1-deficient cells, knockout cells still grew more slowly than untreated controls, indicating that disruption of ncBAF attenuates rather than completely abolishing the therapeutic response. We therefore asked whether this phenotype resulted from an inability to establish the chromatin landscape observed in wild-type cells. ATAC-seq analysis of BRD9-knockout cells demonstrated that the chromatin regions specifically opened by the combinatorial treatment were largely absent in the knockout background, while accessibility changes induced by the individual treatments were also markedly attenuated (Fig. 5d; Extended Data Fig. 6d). In agreement, TF footprinting analysis revealed an almost complete loss of the TF enrichment signature previously identified in treated wild-type cells (Extended Data Fig. 6e). Together, these data demonstrate that BRD9-dependent ncBAF activity is necessary for the full chromatin accessibility landscape that enables activation of the therapy-induced transcriptional regulatory network in NB4 cells.

To determine whether this remodeled chromatin state translated into altered gene expression, we performed RNA-seq on BRD9-knockout cells following treatment. Loss of BRD9 profoundly impaired the transcriptional response, with most genes induced in wild-type cells failing to be activated in the knockout background (Fig. 5e; Extended Data Fig. 6f). Thus, disruption of ncBAF prevents not only the chromatin remodeling events induced by therapy but also their downstream transcriptional output, establishing BRD9 as a key mediator of the differentiation-associated gene expression program in NB4 cells.

Importantly, the requirement for BRD9 was specific to the response elicited by G1-extension. BRD9-deficient NB4 cells retained full sensitivity to retinoic acid treatment, exhibiting growth inhibition comparable to that observed in wild-type cells (Extended Data Fig. 6g). These findings indicate that ncBAF is not broadly required for differentiation signaling but instead specifically couples cell-cycle modulation to chromatin remodeling, activation of the therapy-responsive TF network, and induction of the differentiation-associated transcriptional program.

### LSD1 inhibition in G1-extended susceptible solid tumors halts tumor growth *in vitro* and *in vivo* PDX models

The mechanistic analyses above were performed in AML cell lines. We next asked whether the relationship between cycling speed, p21 expression and sensitivity to LSD1 inhibition generalizes beyond this setting. Therefore, we extended our analysis in two directions: to solid tumor models and to primary patient samples. This arm therefore represents a biological and phenotypic extension rather than a mechanistic re-dissection: the same p21- and cell-cycle–dependent sensitivity relationships established in AML are evaluated here without repeating the epigenomic profiling. Given the growing interest in LSD1 and CDK inhibitors as therapeutic agents across multiple malignancies, we examined melanoma, breast cancer, and small-cell lung carcinoma (SCLC) models — all of which show limited antitumor responses to monotherapy with either LSD1 or CDK inhibitors — to evaluate the broader translational relevance of our findings ^24,84–91^. We first examined *in vitro* solid tumor cell lines, then primary melanoma samples stratified by p21 expression, then the phenotype of drug-tolerant persister (DTP) cells in melanoma and finally validated the combination *in vivo* using melanoma patient-derived xenograft (PDX) models.

In breast cancer, we tested MDA-MB-453 (p21^high^) and MDA-MB-436 (p21^low^), which displayed differential sensitivity to MC2580 corresponding to p21 levels (Fig.6a–c). Notably, MDA-MB-436 cells are RB-deficient and therefore resistant to Palbociclib ^92^: because Palbociclib requires functional RB to enforce G1 arrest, RB-null cells cannot undergo G1 extension via CDK4/6 inhibition. Consistent with this, Palbociclib failed to sensitize MDA-MB-436 cells to LSD1 inhibition (Fig. 6d). However, combined treatment with the HDAC inhibitor TSA and MC2580 significantly inhibited MDA-MB-436 proliferation in a p21-dependent manner (Fig. 6e), demonstrating that pharmacologic extension of G1 length via an alternative route (HDACi-mediated p21 induction) can bypass RB deficiency and sensitize cells to epigenetic therapy. In SCLC — in a cell line previously reported to be sensitive to LSD1 inhibition ^93^ — LSD1 inhibitor response was likewise dependent on p21, as p21 knockdown reversed the anti-proliferative effects (Fig. 6f; Extended Data Fig. 7a–d), confirming the generalizable role of cell cycle in LSD1 inhibitor response across tumor types.

**Figure 6.**
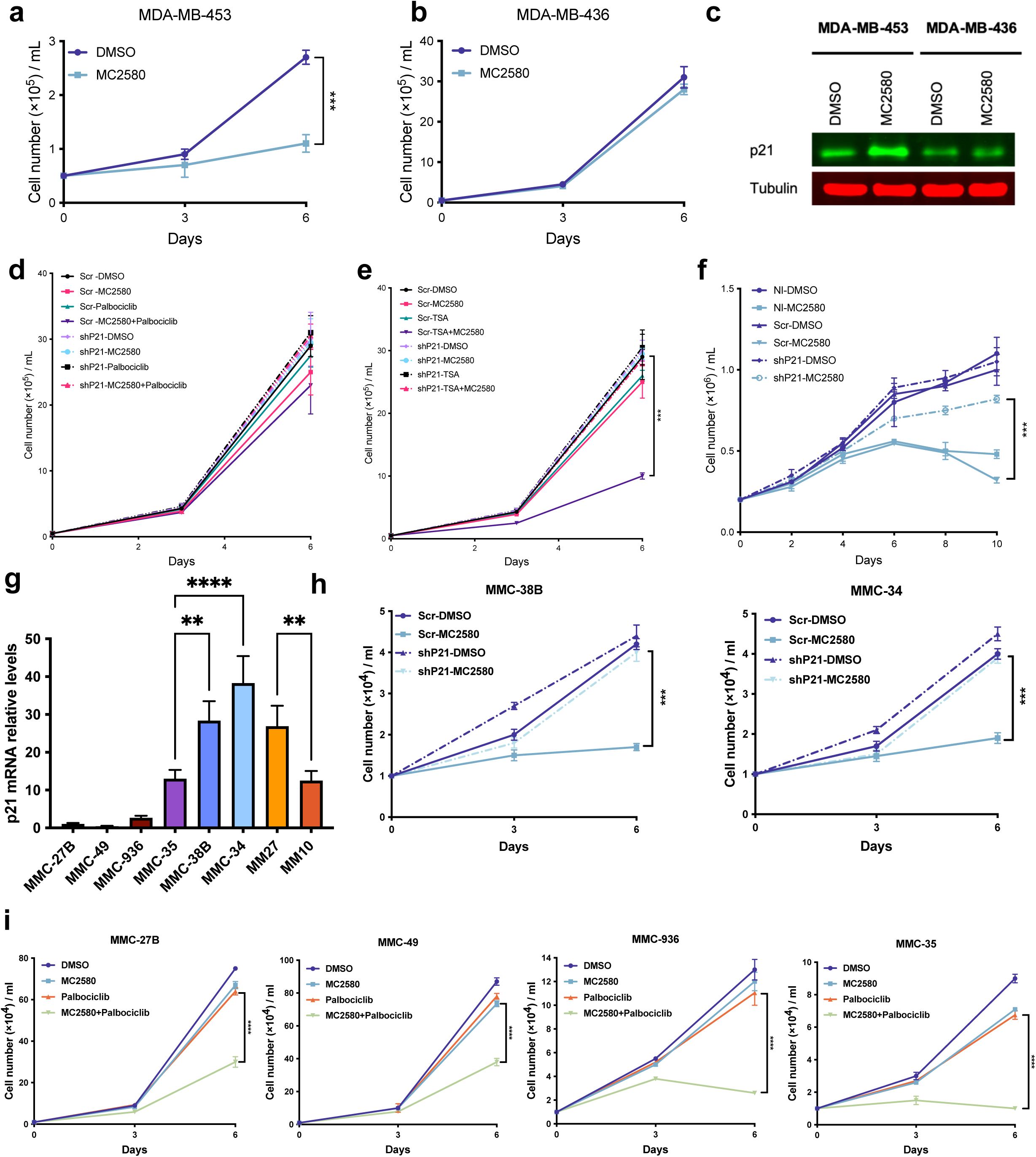
G1 extension sensitizes breast cancer, SCLC and melanoma cells to LSD1 inhibition. **a–b,** Growth curves of breast cancer cell lines treated with DMSO or MC2580 (2 µM). **c,** Western blot of p21 protein in the same breast cancer lines (DMSO vs MC2580, 2 µM). **d–e,** Growth curves of MDA-MB-436 cells with p21 knockdown or Scr control, treated with DMSO, MC2580 (2 µM), Palbociclib (50 nM) (d) or TSA (20 nM) (e), alone or in combination. **f,** Growth curves of NCI-H69 cells with p21 knockdown or Scr control, treated with DMSO or MC2580 (2 µM). **g**, *CDKN1A* mRNA levels in primary melanoma cells and PDX samples, normalized to *GAPDH*. Data are presented as mean ± SD. Statistical significance was assessed by ordinary one-way ANOVA followed by Tukey’s multiple-comparisons test. ns, not significant; ✱P < 0.05, ✱✱P < 0.01, ✱✱✱P < 0.001, ✱✱✱✱P < 0.0001. **h**, Growth curves of MMC-38B and MMC-34 cells stably transduced with the indicated vectors and treated with DMSO or MC2580 (2 µM). **i**, Growth curves of primary melanoma cells treated with DMSO, MC2580 (2 µM), Palbociclib (50 nM), or combinations. Data are mean ± SD of triplicates. All quantitative growth data across the figure were analyzed by one-way ANOVA with Bonferroni’s post-hoc test. P < 0.05 (✱), P < 0.01 (✱ ✱) and P < 0.001 (✱ ✱ ✱).

To validate our findings in more clinically relevant models, we analyzed eight primary patient samples (six for *in vitro* and two for *in vivo* studies; see below), stratified by proliferation rates and p21 expression levels (Fig. 6g). Similar to AML, primary melanoma cells with high p21 levels and slow proliferation were more sensitive to MC2580, while fast-cycling, p21^low^ cells were resistant (Extended Data Fig. 7e, f). p21 knockdown in two p21^high^/MC2580-sensitive primary samples reduced p21 protein levels, prevented its induction upon MC2580 treatment, and increased the S-phase fraction (Extended Data Fig. 7g–l). MC2580 strongly inhibited proliferation and modestly induced apoptosis in control (Scr) cells, while p21 KD rescued cell growth and reduced MC2580-induced apoptosis (Fig. 6h; Extended Data Fig. 7m, n), indicating that p21 mediates LSD1 inhibitor sensitivity in primary melanoma, mirroring findings in AML. Cotreatment with Palbociclib and MC2580 also proved effective in melanoma cells resistant to MC2580 monotherapy, halting proliferation and triggering apoptosis (Fig. 6i; Extended Data Fig. 7o–r).

DTP cells represent a small, reversibly slow-cycling, and drug-resistant population that emerges upon chemotherapy ^94^. Hence, we asked whether chemotherapy-induced selection of slow-cycling cells creates a therapeutic vulnerability to LSD1 inhibition. In melanoma we generated DTPs by treating WM266.4 (naturally p21^high^) and SK-MEL-28 (naturally p21^low^, LSD1i-resistant) melanoma cells with high-dose cisplatin, resulting in 1–4% surviving cells under treatment (Fig. 7a). Both p21 and LSD1 were upregulated in DTPs of WM266.4 cells (naturally higher p21) and SK-MEL-28 cells (naturally lower p21) (Fig. 7b). Pathway analysis of DEGs from single-cell RNA-seq data confirmed that DTPs upregulate pathways related to cell cycle and DNA repair (Extended Data Fig. 7s), and cell cycle profiling showed a clear shift from highly proliferative parental melanoma cells — mainly expressing S and G2/M phase genes — to a slow-cycling state in DTPs, identified by upregulation of a G0/G1 transcriptional profile. The key finding, however, is not simply that DTPs are slow-cycling — a feature intrinsic to their definition — but that they acquire sensitivity to LSD1 inhibition. WM266.4 cells (p21^high^) were eradicated by combining cisplatin with MC2580, with no recoverable viable cells by day 6 of treatment (Fig. 7c). SK-MEL-28 cells (p21^low^, MC2580-resistant) generated DTPs with elevated p21 and LSD1 expression and were likewise eradicated by cisplatin/MC2580 cotreatment (Fig. 7c). Thus, chemotherapy enriches for slow-cycling, drug-resistant cells that are sensitized to LSD1 inhibition.

**Figure 7.**
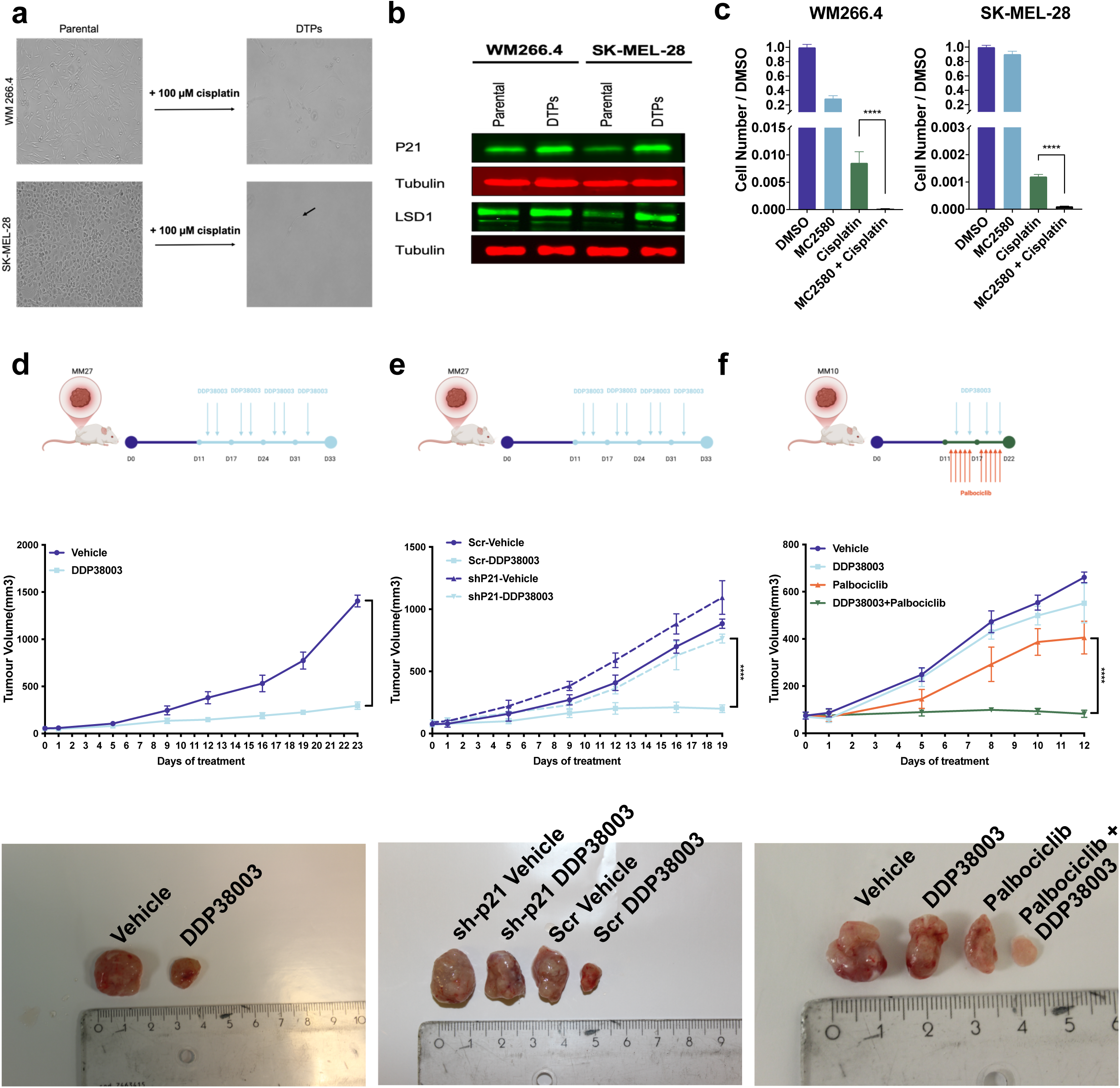
Extended G1 length in melanoma correlates with LSD1 inhibitor sensitivity *in vitro* in DTPs and *in vivo* in PDX models. **a,** Representative images of persister melanoma cells after cisplatin (100 µM, 6 days). **b,** Western blot of parental and DTP melanoma cells for indicated proteins; tubulin loading control. **c**, Relative viability of indicated melanoma cells treated with DMSO, MC2580 (2 µM), cisplatin (100 µM), or combinations (6 days). Mean ± SD of triplicates are shown. **d,** Experimental scheme and tumor growth of MM27 cells in NSG mice treated with vehicle or DDP38003 (experiment workflow created with BioRender.com). **e,** Experimental scheme and tumor growth of MM27 cells transduced with Scr or p21 shRNA in NSG mice treated with vehicle or DDP38003 (experiment workflow created with BioRender.com). **f,** Experimental scheme and tumor growth of MM10 cells in NSG mice treated with vehicle, DDP38003, Palbociclib, or combination. All quantitative growth data across the figure were statistically analyzed using ordinary one-way ANOVA followed by Tukey’s multiple-comparisons test. P < 0.05 (✱), P < 0.01 (✱✱), P < 0.001 (✱✱✱), and P < 0.0001 (✱✱✱✱); ns, not significant. Analysis of the data in panel c was performed on log₂-transformed values.

To assess the *in vivo* efficacy, we selected two melanoma PDX models based on p21 expression: MM27 (p21^high^) and MM10 (p21^low^) (Fig. 6g), with at least 7 mice per group, randomized. As MC2580 is unsuitable for *in vivo* use ^72^, we used its analog DDP38003. DDP38003 dramatically reduced tumor size in MM27 mice (Fig. 7d). p21 knockdown in MM27 tumors accelerated growth and abolished the antitumor effect of DDP38003 (Fig. 7e). In contrast, MM10 tumors, which did not respond to DDP38003 alone, were sensitized by cotreatment with Palbociclib, resulting in reduced tumor burden (Fig. 7f). These data support p21 as a candidate biomarker of LSD1 inhibitor response, a conclusion that will require validation with a defined p21 cutoff and an independent cohort, and demonstrate the efficacy of combining LSD1 inhibition with CDK4/6 inhibition in resistant tumors.

In conclusion, across solid tumor models including melanoma, breast cancer, and SCLC, primary melanoma samples, melanoma DTPs, and *in vivo* PDX models, our findings highlight the therapeutic potential of combining LSD1 inhibition with CDK impairment or p21-inducing agents in tumors with intrinsic or acquired resistance to LSD1 inhibitor monotherapy.

## Discussion

Here, we show that the length of the cell cycle is a critical determinant of the response to epigenetic therapy in tumor cells. Our findings support a conceptual shift in how the cell cycle is viewed in oncology. We propose that the cell cycle is not merely a passive mechanism by which cells execute proliferation in response to growth signals; rather, it functions as a background mechanism of control over cellular fate through the epigenome. By determining the duration of each phase—particularly G1, during which the chromatin landscape is most amenable to remodeling—the cell cycle establishes the conditions under which epigenetic regulators can redirect transcriptional programs. This principle was originally hypothesized in embryonic stem cells, where cell cycle length controls the balance between self-renewal and lineage commitment ^1,2,7,9,11–13,95^. We now show that it operates in tumor cells as well, and that it can be pharmacologically exploited: rather than using cell cycle inhibitors solely to arrest proliferation, low doses can be deployed to extend the epigenetic remodeling window, converting a cytostatic agent into a sensitizer for epigenetic therapy. This constitutes a therapeutic opportunity that is conceptually distinct from conventional anti-mitotic strategies, in which cell cycle inhibitors are dosed at maximum tolerated doses to maximize arrest rather than to tune chromatin accessibility.

In AML, pharmacological extension of G1 by low-dose palbociclib creates a permissive window during which LSD1 inhibition reprograms chromatin on a scale that neither agent achieves alone: in turn, the combination leads to extensive upregulation of genes engaged by myeloid differentiation programs, whereas LSD1 inhibition alone is insufficient and palbociclib alone has minimal transcriptional effect. Mechanistically, LSD1 inhibition installs active histone marks, but the removal of repressive marks—H3K9me3 and H3K27me3—is mainly driven by palbociclib, revealing a double-lock principle in which active mark deposition and repressive mark erasure are separable events gated by cell cycle length and relevant epigenetic targeting. p21 (*CDKN1A*) emerges as the central molecular determinant: tumor cells with high p21 are intrinsically sensitive, whereas p21^low^ cells are resistant but can be sensitized by palbociclib cotreatment. We extend these findings from AML cell lines to solid tumor cell lines (melanoma, breast cancer, SCLC), primary patient melanoma samples stratified by p21, cisplatin-induced drug-tolerant persister cells, and *in vivo* PDX models, demonstrating that the principle generalizes across tumor types and disease settings.

The response to cell cycle prolongation is not necessarily uniform. Each tumor cell carries its own epigenome, shaped by its lineage of origin, driver mutations, and selective history, and will therefore respond differently to G1 extension. Importantly, the nature of the epigenetic vulnerability itself is context-dependent: a given epigenomic state may confer sensitivity not only to LSD1 inhibitors but also to other classes of epigenetic drugs—HDAC inhibitors, EZH2 inhibitors, BET bromodomain inhibitors—depending on which chromatin regulators are rate-limiting for maintaining that state. We postulate, however, that this diversity is not infinite. Rather, we propose that there exist defined sets of epigenomic response—discrete chromatin states that determine which transcriptional programs become accessible upon epigenetic drug treatment and which remain locked. These response classes likely correspond to specific configurations of repressive mark deposition, transcription factor network occupancy, and chromatin remodeling factor availability. Identifying these states prospectively would transform patient stratification from a single-biomarker approach (p21 high versus low) to a multi-dimensional epigenomic classification. To this end, we are performing extensive genetic screens to identify cell context-dependent vulnerabilities that co-determine the epigenomic response to cell cycle prolongation, with the goal of mapping the epigenomic states that predict therapeutic benefit.

Mechanistically, the combination of cell cycle prolongation and epigenetic drug treatment does not simply adjust transcriptional output; it triggers a profound change of cell state, culminating in differentiation. In NB4 cells, the differentiation program induced by palbociclib plus LSD1 inhibition appears molecularly distinct from that triggered by retinoic acid, the canonical differentiation agent in acute promyelocytic leukemia, suggesting that cell cycle– epigenome coupling provides a specific rather than generic differentiation cue. This distinction is further supported by the differential requirement for ncBAF in these two differentiation routes. BRD9 loss impaired differentiation induced by G1 prolongation and LSD1 inhibition, while leaving retinoic acid–induced differentiation largely unaffected, indicating that ncBAF dependency is not a general requirement for differentiation in NB4 cells, but is specifically engaged by the cell cycle–dependent transition. This finding is particularly relevant considering that physiological hematopoietic differentiation relies on stage- and lineage-specific dependencies on chromatin remodeling complexes. Indeed, ncBAF/BRD9 itself is required for selected transitions during normal hematopoietic differentiation ^96^. Our findings therefore raise the possibility that extending G1 does not simply impose a pharmacological differentiation program, but creates a permissive cell-cycle state in which endogenous chromatin-remodeling machinery can execute cell-state transitions. In this model, G1 extension may represent a broadly shared upstream mechanism, whereas the remodeling complexes required to execute the transition are determined by lineage and differentiation state. The differentiation elicited through this route may therefore engage distinct transcription factor networks and chromatin trajectories compared to ligand-induced differentiation, raising the possibility that cell cycle–driven epigenetic reprogramming can access cell fates that conventional differentiation agents cannot. We have opened the way to this concept, but several questions remain to be addressed in future studies: the precise chromatin dynamics that distinguish cell cycle–driven from ligand–driven differentiation, the range of tumor contexts in which this specificity holds, and the identity of the epigenomic states that determine whether differentiation versus alternative cell fates are engaged upon treatment.

The translational implications of this framework are grounded in the recognition that tumors display marked heterogeneity in their proliferative states. In current clinical practice, proliferation is assessed almost exclusively by a single marker, Ki-67, yet this approach is fundamentally limited: Ki-67 is highest in G2 and classifies many proliferating cells in earlier phases as negative, failing to capture the wide range of proliferative states that tumor cells can assume. Recent work using highly multiplexed tissue imaging has shown that only a subset of tumors proliferate with canonical cell cycle dynamics, while others are skewed toward specific phases or exhibit non-canonical combinations of cell cycle regulators, and that this proliferative architecture varies with oncogene activity, spatial location, and therapeutic intervention ^22^. Defining the proliferative state of a tumor will therefore require multivariate markers—combinations of cell cycle regulators such as CDT1, Geminin, phospho-RB, cyclins, and CDK inhibitors—rather than Ki-67 alone. This heterogeneity directly informs therapeutic design. Tumors in a fast-proliferative state—such as AML, where the leukemic blast compartment is characterized by rapid cycling—can be tackled by low doses of cell cycle inhibitors, including palbociclib or next-generation CDK4/6 inhibitors, to extend G1 and sensitize to epigenetic therapy. Conversely, we show that even slow-cycling cells represent a tractable target: cisplatin-induced DTPs, which adopt a reversibly quiescent state with upregulated p21 and LSD1, are eradicated by combining cisplatin with LSD1 inhibition. Thus, both fast- and slow-proliferating tumor compartments can benefit from combinations of cell cycle- and epigenome-targeting drugs, provided that the cell cycle is modulated to create the epigenetic remodeling window. The challenge for clinical translation lies in identifying, for each patient, the epigenomic state that predicts whether cell cycle prolongation will unlock a therapeutic response and in matching that state to the right combination of agents.

Finally, the implications of this work extend beyond cancer to normal physiology. If the cell cycle functions as a background regulator of cellular fate through the epigenome, then the principles we describe—phase duration as a determinant of chromatin remodeling, cell cycle length as a tunable variable for fate redirection—should apply to non-transformed cells as well. Indeed, two features underlying the response observed here are also integral to physiological differentiation: G1 represents a window of increased differentiation competence ^8,97–100^, while progression through normal differentiation trajectories requires stage- and lineage-specific cooperation between transcription factors and chromatin remodeling complexes ^96^. Our findings suggest that these two processes may be functionally coupled: prolonging G1 may establish a temporal window in which the appropriate endogenous remodeling machinery can reshape chromatin, thereby creating the conditions for epigenetic drugs to more effectively redirect cell fate. In this view, the generalizability of the approach would not depend on a specific remodeling complex, epigenetic target, or cancer-associated genetic alteration, but on recapitulating fundamental features of physiological cell-fate transitions—establishing a permissive cell-cycle state and exploiting the context-specific chromatin machinery active within it. This extends the original concept from embryonic stem cells, where cell cycle length controls self-renewal versus differentiation, to a broader biological setting: developing tissues, adult stem cell compartments, and differentiated cells that re-enter the cycle upon injury. By placing the cell cycle back at center stage as an active regulator of epigenetic fate rather than a passive conduit for proliferation signals, we provide a framework that connects stem cell biology, differentiation, and cancer therapy under a unified principle.

## Materials and Methods

### Cell models and genetic tools

#### Cell lines and culture conditions

Cell lines were routinely screened for mycoplasma contamination and authenticated by short tandem repeat (STR) profiling. All cell lines were maintained under standard humidified conditions at 37 °C with 5% CO₂. The AML cell-line panel comprised U-937 (M5), PL-21 (M3; FLT3-ITD), NB4 (M3; PML–RARα), Mono-Mac-6 (M5; KMT2A–MLLT3), KG-1a (AML), MOLM-13 (M5a; KMT2A–MLLT3 and FLT3-ITD), ML-2 (M4; KMT2A–AFDN), GDM-1 (M4), MEG-01 (BCR–ABL1), MV4-11 (KMT2A–AFF1 and FLT3-ITD), UF-1 (M3; PML–RARα), SKNO-1 (M2; RUNX1–RUNX1T1), EOL-1 (FIP1L1–PDGFRA) and Kasumi-1 (M2; RUNX1–RUNX1T1 and KIT-mutant). Unless otherwise specified, cells were cultured in RPMI-1640 supplemented with 10–20% fetal bovine serum (FBS), 2 mM L-glutamine and 1% penicillin– streptomycin. U-937, MOLM-13, ML-2, GDM-1, MEG-01 and EOL-1 were maintained in RPMI-1640 containing 10–20% FBS, whereas PL-21 and UF-1 were maintained in RPMI-1640 containing 20% FBS. KG-1a cells were maintained in Iscove’s modified Dulbecco’s medium (IMDM) supplemented with 20% FBS, and MV4-11 cells were maintained in IMDM supplemented with 10% FBS. Mono-Mac-6 cells were maintained in RPMI-1640 supplemented with 10% FBS, 2 mM L-glutamine, non-essential amino acids, 1 mM sodium pyruvate and 10 µg ml−1 human insulin. SKNO-1 cells were cultured in RPMI-1640 supplemented with 10% FBS and 10 ng ml−1 recombinant human GM-CSF. All media were supplemented with 1% penicillin–streptomycin where indicated, and cultures were maintained in logarithmic growth phase and routinely monitored for mycoplasma contamination. HEK293T cells, used for lentiviral production, were maintained in DMEM with 10% FBS, 2 mM L-glutamine, and 1% penicillin/streptomycin. Solid tumor cell lines, including NCI-H69, SK-MEL-28, WM266.4, MDA-MB-231, MDA-MB-453, and MDA-MB-436, were cultured in media appropriate for each line (IMDM, MEM, DMEM, or DMEM/Ham’s F12) supplemented with 10% FBS, 2 mM L-glutamine, and standard additives (sodium pyruvate and non-essential amino acids where indicated). Primary human melanoma cells and PDX melanoma cells ^101^ were maintained in RPMI-1640 or IMDM, respectively, with 10% FBS, 2 mM L-glutamine, and 1% penicillin/streptomycin.

#### Persister cell derivation and treatment

Persister cells were generated by treating WM266.4 and SK-MEL-28 cells with 100 μM cisplatin for 6 days, refreshing the drug every 48 hours. Unless stated otherwise, pre-derived persister cells were maintained under continuous cisplatin exposure and treated with either DMSO (vehicle) or MC2580. Cell viability was assessed using the CellTiter-Glo assay (Promega) according to the manufacturer’s instructions.

#### Lentiviral vector production and cell transduction

Synthetic oligonucleotides were cloned into vectors containing either a puromycin resistance cassette or GFP reporter for knockdown or overexpression purposes. For knockout experiments, vectors containing both puromycin resistance and GFP were used to express sgRNAs. Vectors were digested with appropriate restriction enzymes, dephosphorylated with Antarctic Phosphatase (New England Biolabs), and purified via 1% agarose gel electrophoresis using the QIAquick Gel Extraction Kit (QIAGEN). Inserts were ligated into vectors at defined molar ratios using T4 DNA Ligase or Quick Ligase (New England Biolabs), and ligation products were transformed into STBL3 competent E. coli via standard heat shock protocols. Transformed colonies were selected on ampicillin-containing LB agar, cultured in LB broth, and plasmid DNA was extracted using QIAprep Miniprep or Maxiprep kits. Insert presence was verified by restriction digest or Sanger sequencing.

HEK293T cells were expanded and transfected with a combination of target plasmid, pCMV-dR8.2, and VSVG using the CaCl₂/HEPES-buffered saline method. Culture medium was replaced 24 h post-transfection. Viral supernatants were harvested, filtered (0.45 µm), concentrated using polyethylene glycol (PEG), and resuspended in PBS for storage at −80 °C. PEG solutions were prepared in MilliQ water with NaCl and filtered under sterile conditions.

Lentiviral shRNA constructs were generated by ligating hairpin-forming oligonucleotides into AgeI/EcoRI-digested pLKO vectors containing an eGFP reporter. The generation of sgRNA-expressing vectors was outsourced to GenScript Biotech (Rijswijk, Netherlands), where sgRNAs were inserted in the ARUNA library vector. Expression plasmids for wild-type and mutant p21 (CDK- or PCNA-binding mutants) were verified by sequencing, and GFP-positive cells were sorted via FACS. FUCCI(CA)2 plasmids containing mCherry and mVenus reporters were obtained from RIKEN BRC (Tsukuba, Japan), and fluorescent cells were similarly isolated by FACS.

Cells were seeded at 5 × 10⁵ per well in 24-well plates. Viral particles, either freshly prepared or concentrated and frozen, were added with 5 µg/mL polybrene. Two to three rounds of transduction were performed as needed, followed by centrifugation at 2500 rpm for 1 h at room temperature and overnight incubation at 37 °C. Successfully transduced cells were selected by antibiotic resistance or fluorescence and isolated via FACS.

### Functional assays

#### Cell proliferation assays

AML cells were seeded at densities of 4 × 10⁵ (UF-1 and Kasumi-1) or 2 × 10⁵ (NB4 and PL-21) per well, and solid tumor cells at optimal growing confluency and treated with the indicated compounds: 2 µM MC2580, 50/40 nM palbociclib, 20 nM SAHA, 20 nM TSA, 1 µM UNC1999, 500 nM Milciclib, and 1 µM DDP38003. Note that the concentrations of SAHA and TSA were adjusted to minimize antiproliferative effects, yet maintaining the ability to induce p21 expression. Proliferation was assessed in triplicate. Cell viability was determined by Trypan blue exclusion (1:1, Sigma) using a Bio-Rad TC20™ automated cell counter or a hemocytometer. For growth-curve experiments of sgRNA-expressing cells, cells were counted by flow cytometry using a BD FACSCelesta (BD Biosciences) equipped with a High Throughput Sampler (HTS). For each sample, a fixed acquisition volume of 50 µL was analyzed. Samples were diluted 1:2 prior to acquisition. At each time point, the number of mCherry⁺/GFP⁺ cells was determined by flow cytometry, corresponding to cells expressing Cas9 and sgRNA, respectively, and cell numbers were calculated as cells/mL by multiplying the number of double-positive events detected in the 50-µL sample by 40. At time zero, cells were seeded at 2 × 10⁵ cells/mL in all conditions, and the number of mCherry⁺/GFP⁺ cells was estimated from the proportion of double-positive cells measured in the pre-plating cell suspension. Thus, the initial number of double-positive cells was calculated as 2 × 10⁵ × the fraction of mCherry⁺/GFP⁺ cells. This correction accounted for differences in the proportion of Cas9/sgRNA-positive cells between samples at the time of seeding. Competitive cell growth following treatment of knockout cells was assessed by co-culturing NB4 iCas9 cells (GFP^−^) with sgRNA-expressing cells (GFP^+^) at a 1:1 ratio. 0.5 µg/mL doxycycline was maintained during knockout-cell proliferation to preserve inducible Cas9 expression. Absolute numbers of GFP^−^ and GFP^+^ cells were determined by flow cytometry using a BD FACSCelesta (BD Biosciences) equipped with a High Throughput Sampler (HTS). For each sample, a fixed acquisition volume of 50 µL was analyzed, and data were processed using FlowJo software (BD Biosciences).

#### Colony-forming unit (CFU) assay

Cells (5 × 10³) were plated in triplicate in methylcellulose medium (MethoCult™ GF H4435 Enriched; StemCell Technologies) supplemented with either DMSO or MC2580 (2 µM). For serial replating, cells recovered from colonies were reseeded in fresh semi-solid medium of the same composition. Colonies were scored every 7–10 days after seeding.

#### Morphological analysis

Cells were deposited onto glass slides using a Cytospin™ 4 cytocentrifuge and stained with May-Grünwald-Giemsa. Slides were first incubated in May-Grünwald solution (Sigma Aldrich) for 8 min, washed six times with deionized water, and then incubated in Giemsa solution diluted 1:19 with distilled water for 30 min. After three additional washes in distilled water, slides were air-dried and coverslipped using Eukitt® mounting medium for long-term preservation.

#### Apoptosis analysis by Annexin V/Propidium Iodide staining

Cells were treated as indicated, collected, washed with PBS, and counted. For each sample, 5 × 10⁵ cells were resuspended in 50 µL Annexin V binding buffer (10 mM HEPES, 150 mM NaCl, 1 mM MgCl₂, 3.6 mM CaCl₂, 5 mM KCl) and labeled with Annexin V-FITC for 1 h at room temperature. Cells were then diluted in 500 µL PBS with 1 µL propidium iodide (50 µg/mL) and analyzed by flow cytometry (BD FACSCelesta, BD Biosciences). Annexin V-FITC detects phosphatidylserine exposure, whereas propidium iodide identifies late-apoptotic and necrotic (membrane-permeable) cells.

### Screening

#### Drug screening

##### Compound library

The compound library used for screening consisted of 290 small molecules, including FDA-approved drugs (Selleck L1900 epigenetics compound library), provided as 10 mM stock solutions in dimethyl sulfoxide (DMSO) and stored at −80 °C. Compounds were subsequently diluted to 1 mM and maintained at −20 °C prior to use. Working dilutions were freshly prepared in assay medium immediately before treatment, maintaining a final DMSO concentration of 0.23% (v/v) in all experimental conditions.

##### Drug screening assay

NB4 cells were seeded at 4 × 10⁵ cells/mL and treated with 50 nM palbociclib; parallel samples were treated with vehicle (DMSO) as control. Pre-treated cells were then plated at 250 cells per well in white 96-well plates (Thermo Scientific Nunc, catalog no. 136102) and screened in duplicate with three concentrations (2300, 233, and 2.33 nM) from the compound library in medium containing 50 nM palbociclib. Cell seeding and compound dispensing were performed using a liquid-handling robot (Tecan Fluent).

Treatments were performed for six days, and cell viability was determined using the CellTiter-Glo luminescent assay (Promega). The luminescence was measured with a Spark microplate reader (Tecan) using 30ms per well of integration time. Cells treated with DMSO alone or with 50 nM palbociclib alone were included as controls.

##### Data Normalization and Hit Identification

Luminescence values measured at the end of treatment were normalized to Palbociclib + vehicle-treated control wells (DMSO), which were used as negative control. Normalization was performed by calculating the ratio between the luminescence signal of each treated sample and the mean luminescence signal of the corresponding vehicle control wells. The resulting normalized values were expressed as fold change relative to the vehicle control.

The formula is:

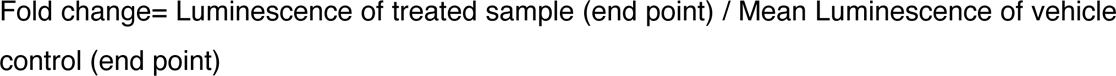

Hit identification was defined as a fold change of < 30% (for at least one replicate of the compound) since the administrated dose was an average IC_70_ of the library. Drug targets were assigned using the Selleck library annotation and manually curated target information. Multi-target drugs were expanded into individual drug–target pairs, and for each target we counted the number of hit compounds and the total number of screened compounds targeting it. Non-hit counts were calculated as total minus hit compounds per target.

Statistical Analysis and Quality Control parameters: Z’ factor has been used to determine assay dynamic range and data variation using both the positive and negative controls on each plate. A measure of statistical effect size and calculated as the means (µ) and standard deviation (σ) of both positive (p) and negative (n) controls: Z′ factor = 1 − 3(σσ_p_+σσ_n_)/|μμ_p_-μμ_n_|. Acceptable Z′-factor values range between 0.5 and 1; a plate is invalidated if the Z′-factor falls below 0.3 ^102^. Corresponding values can be provided upon request.

#### CRISPR-Cas9 screening

The doxycycline-inducible Lenti-iCas9-neo vector (Flag-iCas9-P2A-GFP; Addgene plasmid #85400; RRID:Addgene_85400), originally described by Cao et al., was obtained from Addgene ^103^. The GFP reporter was replaced with mCherry and the P2A sequence was replaced with T2A, generating the Flag-iCas9-T2A-mCherry construct used in this study. NB4 cells were infected with an inducible Cas9-T2A-mCherry expression vector, and selected with 400 µg/ml G418 for one week. Selected cells were plated at low density in cytokine-free semi-solid methylcellulose medium (MethoCult™ H4230, STEMCELL Technologies) to allow isolation of single-cell clones. After two weeks, individual colonies were transferred to liquid culture and subsequently expanded. Colonies were screened by testing the efficiency of different sgRNAs using the ICE algorithm developed by Synthego ^104,105^.

The ARUNA library is a subset of the GecKO library ^106^. It targets 652 chromatin-related genes with 6 sgRNAs targeting each gene, and 60 non-targeting sgRNAs were included as controls.

The screening was performed in biological triplicate, maintaining a 500× representation of the library throughout every step. NB4-iCas9 cells were infected with the ARUNA library at low MOI (<0.3) calculated 72 h post-infection by GFP positivity. Cells were then selected for 48 h with 2 µg/mL puromycin. Puromycin-resistant cells were treated with 0.5 µg/mL doxycycline to induce Cas9-mCherry expression. A minimum of 4 million cells/sample were collected 48 h after doxycycline exposure as T0 of the experiment. Samples were then divided into vehicle (DMSO) or treated (palbociclib 50 nM) and maintained for 14 days.

Genomic DNA from T0 and T14 of vehicle/Palbociclib-treated cells was extracted using the DNeasy Blood and Tissue Kit (Qiagen). sgRNA sequences were amplified by two rounds of PCR, compatible with Illumina sequencing. The resulting libraries were sequenced (2 × 61 bp paired-end) on an Illumina NovaSeq 6000 sequencer.

### Flow cytometry, sorting, and imaging

#### Flow cytometry analysis

FUCCI(CA)2-expressing cells (∼1 × 10⁶ per sample) were harvested, washed with 1% BSA in PBS, and fixed by sequential incubation in PBS with 2% formaldehyde (20 min on ice) followed by dropwise addition of ethanol to a final concentration of 75%, with at least 30 min on ice. Fixed cells were stored at 4 °C as needed. For DNA staining, cells were resuspended in DAPI (5 µg/mL in PBS) and incubated overnight at 4 °C. Cell-cycle, FUCCI(CA)2 reporter, and mCherry and mVenus expression analyses were performed by flow cytometry (BD FACSCelesta, BD Biosciences, Oxford, UK), with unstained cells included as negative controls.

#### Fluorescence-activated cell sorting of live or fixed cells

Cells (≥5 × 10⁵ per sample) were resuspended in sorting solution (PBS with 2% FBS, 1% penicillin/streptomycin, 0.3% Gentamicin, and 1X Amphotericin B) at 5 × 10⁶–1 × 10⁷ cells/mL and filtered through a 70 µm mesh to remove aggregates. Both infected and uninfected controls were processed. Cells were collected into medium containing 33% FBS, 3% penicillin/streptomycin, and 0.3% Gentamicin. Post-sorting, cells were maintained in standard culture medium with 1X Amphotericin B until downstream use or cryopreservation. Sorting was performed on a FACSAria™ or FACSMelody™ cell sorter (BD Biosciences, Oxford, UK). Identification of cells in G1 phase was performed by FUCCI(CA)2 (mCherry^+^/mVenus^-^ cells). For BRD9-KO cells, cells were stained with 10 µg/mL Hoechst 33342 for 90 min prior to acquisition. The two staining methods for G1 identification were cross-checked to ensure reproducibility of the G1-sorted population ^28^.

#### Live-cell imaging and cell cycle quantification

AML cells (2.5 × 10^5 per well) were seeded in triplicate into Tethis (Milan, Italy) SBS 12-well plates and processed following ^28^. In brief, cells were treated with the indicated palbociclib doses in MethoCult™ H4230 RPMI-1640 (80 mL) and RPMI-1640 (20 mL) supplemented with 10% FBS, 2 mM glutamine, and 1% penicillin/streptomycin.

Time-lapse imaging was performed using a Leica Thunder Imager equipped with a Lumencor Spectra X Light Engine, motorized stage, and Leica DFC9000 GTC camera. Images were acquired with a 20×/0.75 NA air objective, 2 × 2 binning, and LAS X software (v3.7.5.24914). Fluorescence from mCherry and mVenus was recorded using excitation filters of 540–580 nm and 460–500 nm, dichroic mirrors at 585 and 505 nm, and emission filters of 592–668 nm and 512–542 nm, respectively. A brightfield channel was acquired in parallel. Per well, 20–25 fields of view were imaged, with manual focus calibration and adaptive focus control to ensure stability. Time-lapse imaging was performed for 72 h, with acquisitions every 60 min to minimize phototoxicity ^28^.

Imaging data were imported into R v4.3 for analysis. Cycling cells were identified using an in-house machine-learning pipeline developed with tidymodels v1.1.0 and timetk v2.8.3. Quantification of cell cycle phases was performed using tidyverse v2.0.0 ^28^.

### Molecular and proteomic assays

#### RNA extraction and quantitative PCR (qPCR)

Total RNA was extracted from 2–4 × 10⁶ cells using 1 mL TRIzol reagent (Invitrogen) or equivalent lysis buffer. Cell pellets were resuspended and homogenized by pipetting, and lysates were processed with the RNeasy Mini Kit (Qiagen) or, when applicable, the Quick-RNA Miniprep Kit (Zymo Research), according to the manufacturers’ instructions. RNA concentration and purity were determined using a NanoDrop ND-1000 spectrophotometer or Qubit DNA HS assay (Thermo Fisher Scientific).

Complementary DNA (cDNA) was synthesized from purified RNA using OneScript Plus Reverse Transcriptase (ABM) or the SuperScript II Kit (Invitrogen), following the manufacturers’ protocols.

#### Immunoblotting

Cell pellets were lysed in 100–200 µL of SDS lysis buffer (2% SDS, 10% glycerol, 50 mM Tris-HCl) or 8 M urea buffer, supplemented with a protease inhibitor cocktail (Sigma-Aldrich, Cat. no. 11836170001). Lysates were centrifuged at 13,000 rpm for 15 min at 4 °C, and protein concentration was determined using the Bradford assay (Bio-Rad). Proteins (50–80 µg) were mixed with Laemmli buffer (containing β-mercaptoethanol and bromophenol blue), denatured at 95 °C for 10 min, and resolved by SDS–PAGE. Proteins were transferred to nitrocellulose membranes (Whatman) in transfer buffer (1×, 20% methanol) at 100 V for 1 h at 4 °C.

Membranes were blocked in TBS-T (0.1% Tween-20, 5% non-fat dry milk) and incubated with primary antibodies (see Table of reagents, antibodies, and supplies) diluted in blocking solution for 1 h at room temperature or overnight at 4 °C. After three washes in TBS-T (10 min each), membranes were incubated with IRDye near-infrared (LI-COR) or other relevant secondary antibodies diluted in blocking solution for 30–60 min at room temperature. Following three additional washes, signals were detected using the Odyssey imaging system (LI-COR) or by enhanced chemiluminescence (ECL) and a Bio-Rad Gel-Doc XR molecular imager.

#### Mass spectrometry of hPTMs

Approximately 4 µg of histone octamers were mixed with isotopically labeled histones in equimolar amounts as internal standards and separated by 17% SDS-PAGE ^41^. Excised histone bands were chemically acylated with propionic anhydride, digested in-gel with trypsin, and subjected to N-terminal derivatization with phenyl isocyanate (PIC) ^40^. Peptides were separated by reversed-phase chromatography on a 25-cm EASY-Spray column (75 µm ID, PepMap C18, 2 µm; Thermo Fisher Scientific) coupled online to a Q Exactive Plus mass spectrometer via an EASY-Spray™ Ion Source ^40^.

RAW data were processed with EpiProfile 2.0 and manually curated for quality control ^107^. Quantification of histone modifications was expressed as relative abundance (%RA), calculated as the area under the curve (AUC) of each modified peptide normalized to the sum of all forms of that peptide × 100. Ratios of %RA between light (sample, L) and heavy (internal standard, H) channels (L/H) were then determined.

### Genomic assays

#### ATAC-seq

Cells were freshly isolated from *in vitro* culture and FACS-sorted. All procedures before transposition were carried out on ice using pre-chilled buffers and with minimal handling time to preserve chromatin integrity. Cells were washed with 1× PBS, counted, and 5 × 10^4^ cells per sample were pelleted at 500 × g for 10 min at 4 °C using a swinging-bucket rotor. After careful removal of the supernatant, cells were resuspended by three pipetting cycles in 50 µl ice-cold ATAC resuspension buffer (ATAC-RSB; 10 mM Tris-HCl, pH 7.4, 10 mM NaCl and 3 mM MgCl2) supplemented with 0.1% NP-40, 0.1% Tween-20 and 0.001% digitonin and incubated on ice for 3 min. Lysis was stopped by adding 1 ml ice-cold ATAC-RSB supplemented with 0.1% Tween-20 but without NP-40 or digitonin. Samples were mixed by three gentle inversions and nuclei were pelleted at 500 × g for 10 min at 4 °C.

The supernatant was carefully removed and nuclei were resuspended by six pipetting cycles in 50 µl transposition mixture containing 10 µl 5× Tn5 buffer (50 mM Tris-HCl, pH 8.4, and 25 mM MgCl2), 1 µl Tn5 transposase, 16.5 µl 1× PBS, 0.25 µl 2% digitonin, 0.5 µl 10% Tween-20 and 21.75 µl nuclease-free water. Samples were briefly centrifuged and incubated at 37 °C for 30 min in a thermomixer with mixing at 600 r.p.m. The reaction was terminated by adding 20 µl cleanup solution containing 10 µl cleanup buffer (900 mM NaCl and 30 mM EDTA), 4 µl 5% SDS, 2 µl proteinase K (20 mg ml−1) and 4 µl water, followed by incubation at 40 °C for 30 min. Transposed DNA was purified using Agencourt AMPure XP SPRI beads (Beckman Coulter) at a 1.5× bead-to-sample ratio. Beads were equilibrated to room temperature for 30 min, vortexed thoroughly, and 105 µl beads was added to each sample. Samples were mixed by pipetting 15 times, incubated for 5 min at room temperature and placed on a magnetic rack for 5 min. After removal of the supernatant, beads were washed twice with 100 µl freshly prepared 80% ethanol and air-dried until residual ethanol had evaporated. DNA was eluted in 22 µl EB buffer (10 mM Tris-HCl, pH 8.0) or molecular-grade water by pipetting 25 times and incubating for 5 min at room temperature. After magnetic separation, 20 µl eluate was transferred to a fresh tube.

Libraries were generated by PCR using sample-specific indexed i5 and i7 primers. For each reaction, 18 µl transposed DNA was combined with 2 µl i5 primer (10 µM), 2 µl i7 primer (10 µM), 8.8 µl 5× KAPA buffer, 1.32 µl 10 mM dNTPs, 0.88 µl KAPA HiFi polymerase and 11 µl water, for a final volume of 44 µl. The amplification master mix was prepared without DNA or primers, with polymerase added last, and activated at 93 °C for 3 min. PCR was performed at 72 °C for 5 min and 98 °C for 2 min, followed by 15 cycles of 98 °C for 20 s, 63 °C for 30 s and 72 °C for 1 min, with a final hold at 4 °C.

Amplified libraries were adjusted to 50 µl with 6 µl molecular-grade water and size-selected using SPRI beads. First, 32.5 µl beads (0.65×) was added, samples were mixed and incubated for 5 min at room temperature, and beads were magnetically separated for 4 min. The supernatant, containing fragments <500 bp, was transferred to a fresh tube. To remove residual primers, 57.5 µl fresh SPRI beads was added to the recovered supernatant. After mixing and incubation for 4 min at room temperature, beads were magnetically separated, washed twice with 100 µl 80% ethanol and air-dried for 4 min. Libraries were eluted in 22 µl molecular-grade water, incubated for 2 min at room temperature and magnetically separated for 4 min, after which 20 µl eluate was transferred to a fresh tube and stored at −20 °C.

Library concentration was measured using a GloMax instrument and fragment-size distributions were assessed using a TapeStation with HS D5000 ScreenTape. Libraries were expected to show prominent fragment populations at approximately 180–200 bp and 310– 330 bp; libraries containing higher-molecular-weight fragments or residual primers were subjected to additional size selection or SPRI cleanup, respectively. Libraries could initially be evaluated by low-depth sequencing (∼5 million reads per sample), with approximately 10– 20 million genome-aligned reads per sample used as the target for adequate coverage. Library quality was further evaluated from the proportion of mitochondrial reads and PCR duplicates ^108^.

#### ChIP-seq

Chromatin immunoprecipitation (ChIP) was performed using an optimized low-input protocol, with all conditions processed in biological duplicate. Approximately up to 15 × 10^6^ cells processed per condition to obtain approximately 200 µg chromatin per IP. Control and treated samples were adjusted to identical cell numbers and volumes before fixation. Cells were crosslinked by addition of 37% formaldehyde to a final concentration of 1% and incubated for 10 min at room temperature with agitation. Crosslinking was quenched with 2 M glycine at a final concentration of 0.125 M for 5 min at room temperature. Cells were subsequently pelleted at 4 °C and washed twice with ice-cold PBS.

Cell pellets were lysed in 200 µl SDS lysis buffer (50 mM Tris-HCl, pH 8.1, 0.5% SDS, 100 mM NaCl, 5 mM EDTA and 0.02% NaN3) supplemented immediately before use with cOmplete Mini protease inhibitor cocktail (Roche). Lysates were incubated for 2 min at room temperature and processed immediately whenever possible. For sonication, 100 µl Triton dilution buffer (100 mM Tris-HCl, pH 8.6, 5% Triton X-100, 100 mM NaCl and 5 mM EDTA), supplemented with protease inhibitors, was added to each 200-µl lysate. Samples were maintained on ice and sonicated in 1.5-ml Bioruptor Pico microtubes using a Bioruptor Pico (Diagenode) maintained at 4 °C. Chromatin was fragmented using 46 cycles of 30 s sonication followed by 30 s rest, with gentle pipette mixing after the first 23 cycles, to obtain fragments predominantly between 100 and 300 bp.

Chromatin fragmentation was verified using an aliquot of each sample. Briefly, 60 µl sonicated chromatin was combined with 40 µl solution containing 10 µl 10% SDS, 10 µl 1 M NaHCO3 and 20 µl water and incubated at 65 °C for 1 h. Samples were treated sequentially with 2 µl RNase A for 15 min at room temperature and 3 µl proteinase K for 30 min at 65 °C. DNA was purified using a Qiagen PCR purification kit, eluted in 15 µl elution buffer and analysed on a 1% agarose gel to confirm a fragment distribution of approximately 100–300 bp.

Before immunoprecipitation, sonicated chromatin was diluted to 1 ml with IP buffer supplemented with protease inhibitors and pre-cleared for 2 h at 4 °C with BSA-blocked Protein A Sepharose beads (20 µl per 10^7^ cells) under rotation. Protein A Sepharose beads were blocked by washing three times in TE buffer, resuspending as a 50% slurry and incubating with lipid-free BSA at 0.5 mg ml−1 for 2 h to overnight at 4 °C, followed by three additional TE washes. After pre-clearing, beads were removed by centrifugation and the clarified chromatin was recovered.

Chromatin concentration was determined by Bradford assay at 595 nm, and the amount required to obtain a fixed amount of chromatin per IP (ranging 200-1000 µg) across each experiment was calculated depending on estimated target’s abundance. For each IP, chromatin was adjusted to 1 ml, while 25 µl (2.5%) was retained as input. Primary antibody was added at approximately 0.5–2 µg per IP for histone modifications or 3-10 µg per IP for transcription factors (please refer to Table 1); matched rabbit or mouse IgG was used at the same concentration for mock controls. Thirty microlitres of Protein A or Protein G magnetic beads, selected according to the antibody isotype, was added to each IP, and samples were incubated overnight at 4 °C with rotation.

Immune complexes were collected magnetically and washed on ice with 1 ml cold buffer per wash. Beads were washed three times with Mixed Micelle buffer (5.2% sucrose, 1% Triton X-100, 150 mM NaCl, 0.2% SDS, 20 mM Tris-HCl, pH 8.1, 5 mM EDTA and 0.02% NaN3), three times with Buffer 500 (500 mM NaCl, 1% Triton X-100, 50 mM HEPES, pH 7.5, 0.1% sodium deoxycholate, 1 mM EDTA and 0.02% NaN3), three times with LiCl detergent buffer (250 mM LiCl, 0.5% sodium deoxycholate, 0.5% NP-40, 1 mM EDTA, 10 mM Tris-HCl, pH 8.0, and 0.02% NaN3), and finally twice with TE buffer. During each wash, beads were resuspended and magnetically recovered; the final wash with each buffer was incubated for 5 min at 4 °C with rotation.

Chromatin was eluted and crosslinks were reversed by resuspending IP beads in 200 µl decrosslinking solution (0.1 M NaHCO3 and 1% SDS). Input samples were brought to the same final volume by addition of 175 µl decrosslinking solution to the retained 25 µl input. IP and input samples were incubated at 65 °C in a thermomixer at 1,200 r.p.m., followed by addition of 3 µl proteinase K and overnight incubation at 65 °C. Samples were briefly centrifuged, IP beads were magnetically separated, and the supernatants were purified using a Qiagen PCR purification kit by addition of five sample volumes of PB buffer and processing according to the manufacturer’s instructions. DNA was eluted in a total volume of 70 µl EB buffer and quantified using the Qubit DNA High Sensitivity assay. Purified ChIP and input DNA was subsequently used for qPCR validation and/or ChIP-seq library preparation.

#### RNA-seq

Total RNA was extracted and quality-checked using the Agilent Bioanalyzer with a picoRNA chip. Libraries were prepared from 0.1–1 μg of RNA using the Illumina TruSeq v.2 RNA Sample Preparation Kit, following the manufacturer’s instructions. Polyadenylated mRNA was enriched through two rounds of capture using oligo-dT magnetic beads under denaturing conditions. RNA was fragmented using divalent cations at elevated temperature, and first-strand cDNA synthesis was performed with random hexamers and SuperScript II reverse transcriptase (Invitrogen). Second-strand synthesis was carried out with DNA Polymerase I and RNase H.

cDNA was purified using AMPure XP beads, end-repaired, 3′-adenylated, and ligated with Illumina-specific adapters. Libraries were amplified by 15 cycles of PCR using Illumina primer mixes. Final library quality and concentration were verified using an Agilent Bioanalyzer 2100 with a high-sensitivity DNA assay. Sequencing was conducted on an Illumina platform according to standard protocols for poly(A)-selected RNA.

### In vivo

DDP38003 was prepared in a vehicle of 40% PEG-400 in 5% glucose solution, while palbociclib was dissolved in sterile-filtered 50 mM sodium lactate, pH 4 (Sigma). A total of 2.5 × 10⁵ patient-derived melanoma cells were subcutaneously injected into 6–8-week-old NSG mice in Matrigel. Tumor volume was calculated using the modified ellipsoid formula: ½ × (length × width²). Treatment commenced when tumors reached ∼50–100 mm³, with DDP38003 administered orally at 16.8 mg/kg twice weekly and palbociclib at 75 mg/kg five times per week. For combination treatments, at least 4 h separated LSD1 and CDK4/6 inhibitor dosing, with palbociclib given first when both were administered on the same day. All *in vivo* procedures were approved by the institutional animal welfare committee (OPBA), notified to the Ministry of Health in accordance with Italian law (IACUC No. 758/2015), and performed following EU Directive 2010/63. The project number authorized by the Ministry of Health, Italy is 139/2023-PR.

### Computational analyses

#### Data and statistical analyses

##### ChIP-seq analysis

Raw paired-end ChIP-seq reads were quality-filtered and aligned to the hg18 reference genome using the nf-core/chipseq v2.0.0 pipeline with Bowtie2 as the aligner ^109,110^. PCR duplicates were removed, and blacklisted genomic regions were excluded. Peak calling was performed with MACS2 for H3K27ac, H3K4me3, and H3K27me3, whereas H3K9me3 enrichment domains and BRD9-enriched regions were identified using SICER2 (v1.0.2) ^111,112^. Genomic annotation was performed using ChIPseeker v1.32.1, and enrichment heatmaps were generated with deepTools v3.5.1 ^113,114^.

##### ATAC-seq analysis

Raw paired-end ATAC-seq reads were aligned to the hg18 reference genome using the nf-core/atacseq v2.1.2, with Bowtie2 as the aligner and MACS2 for peak calling ^110,111^. Peak calling was performed with MACS2 using an FDR threshold of 0.001 (--macs_fdr 0.001), and consensus peaks were defined as peaks detected in at least 3 of 5 biological replicates (--min_reps_consensus 3). Peak overlap and treatment-specific or shared accessible regions were identified using the findOverlapsOfPeaks function from ChIPpeakAnno ^115^. Motif discovery was performed using GimmeMotifs v0.18.0 ^116^.

Differential transcription factor footprinting analysis was performed using TOBIAS v0.15.1 with the ATACorrect, ScoreBigWig, and BINDetect modules ^74^. Each treatment was compared with the corresponding DMSO control using consensus peak sets. The same workflow was applied to treatment-specific and shared accessible regions by providing the corresponding regions through the --regions option in ScoreBigWig and the --peaks option in BINDetect. For each treatment versus DMSO comparison, the top 20 positively and negatively differentially bound TF motifs were selected based on the BINDetect p-value for downstream analyses. Tornado plots were generated using deepTools v3.5.1 ^114^.

#### RNA-seq analysis

Reads were generated as follows: NB4 WT, NB4 BRD9 KO and Kasumi (150-bp paired-end), PL-21 (51-bp paired-end), NB4 and UF1 (51-bp paired-end). Reads were aligned to the hg18 reference genome using nf-core/rnaseq v3.9, with STAR as the aligner and Salmon for transcript quantification ^117,118^. Differential expression analysis was performed with tidybulk (v1.8), employing edgeR quasi-likelihood methods and TMM normalization. Differentially expressed genes (DEGs) were defined as those with *FDR* ≤ 0.01 and an absolute log fold change ≥ 1.5) ^119,120^.

Pathway and upstream regulator analyses were performed using QIAGEN IPA (QIAGEN Inc., https://digitalinsights.qiagen.com/IPA) ^121^, while over-representation analysis (ORA) was conducted with EnrichR ^122^ (WikiPathways, Reactome, KEGG).

#### Single-cell RNA-seq analysis

Demultiplexing was performed using 10x Genomics Cell Ranger (mkfastq) with default parameters, and transcript quantification was carried out with Cell Ranger v2.1.1 count against the hg19 reference genome. Quality control and filtering were conducted in Seurat v4 ^119^, excluding cells with fewer than 200 or more than 2,500 detected features, or with >5% mitochondrial reads. After filtering, 2,180 and 4,409 cells were retained for the two DMSO replicates and 1,654 and 2,650 cells for the 6-day treatment replicates.

All samples were integrated using the IntegrateData function in Seurat, followed by scaling. UMAP dimensionality reduction was performed using the first 50 principal components. Unsupervised clustering was carried out using the first 50 principal components with a resolution parameter of 0.5.

Pathway enrichment and upstream regulator analyses were performed on the 6-day treatment samples (log₂ fold change > |1.5| versus DMSO) using (QIAGEN Inc., https://digitalinsights.qiagen.com/IPA) ^121^.

#### CRISPR-Cas9 screening analysis

Raw reads (61 bp) were trimmed using seqtk trimfq (v 1.3) (parameters: -b 29 -e 18) in order to retain only the variable barcode sequence. The library fasta file was indexed with bowtie-build ^110^. Trimmed reads were then aligned to the ARUNA library using bowtie (v 1.2.2) with the following parameters: bowtie -v 0 -p 6 -a -m 1 --norc $library -q $Sample.fastq -S ^110^.

Alignment files were processed with the samtools suite (view, sort, index, idxstats v 1.12) to obtain counts of mapped read segments for each sample, replicate, and time point, which were subsequently used for downstream analysis and visualization ^123^.

Gene-level statistics were calculated using MAGeCK (mageck test -k), performing paired comparisons between treatment (in triplicate) and DMSO ^124^. From the resulting gene_summary.txt files, the top 30 positively and negatively enriched genes (excluding non-targeting controls) were extracted based on positive or negative ranking scores. Relative log fold-change (negLFC/posLFC) and FDR values (negFDR/posFDR) were visualized with scatterplots generated using custom R scripts. Genes identified as significantly enriched or depleted were considered positive or negative hits, respectively, and were carried forward for further analysis.

#### Use of generative artificial intelligence

ChatGPT (OpenAI) was used as a supplementary tool during manuscript preparation. Its use was limited to supporting the synthesis and comparison of relevant literature and to improving the clarity, structure, and language of selected sections of the text. Any AI-assisted output was critically reviewed by the authors and, where applicable, verified against the original scientific sources. ChatGPT was not used for the generation, acquisition, processing, analysis, or modification of experimental data, nor was it used to draw scientific conclusions. All scientific interpretations, methodological decisions, and final written content remained the responsibility of the authors.

#### Reagents and resources

**Table 1.**
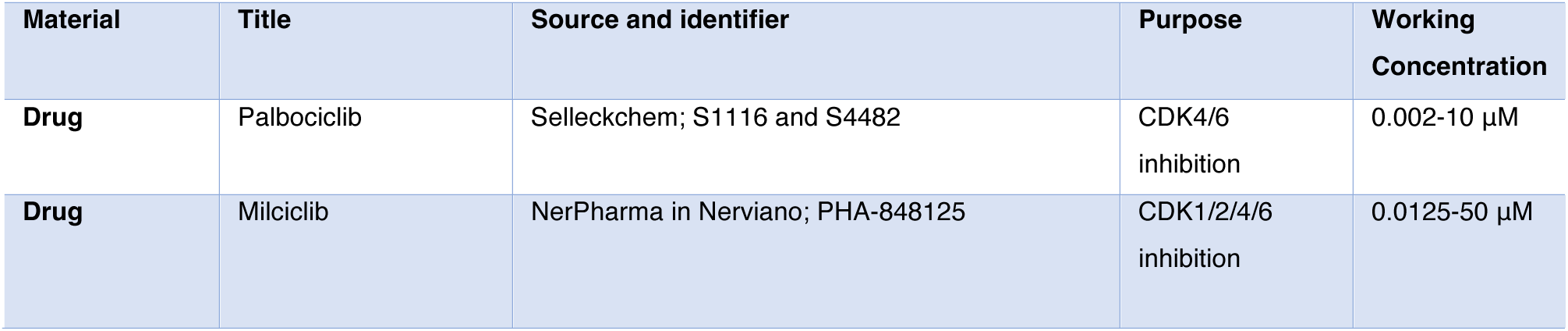
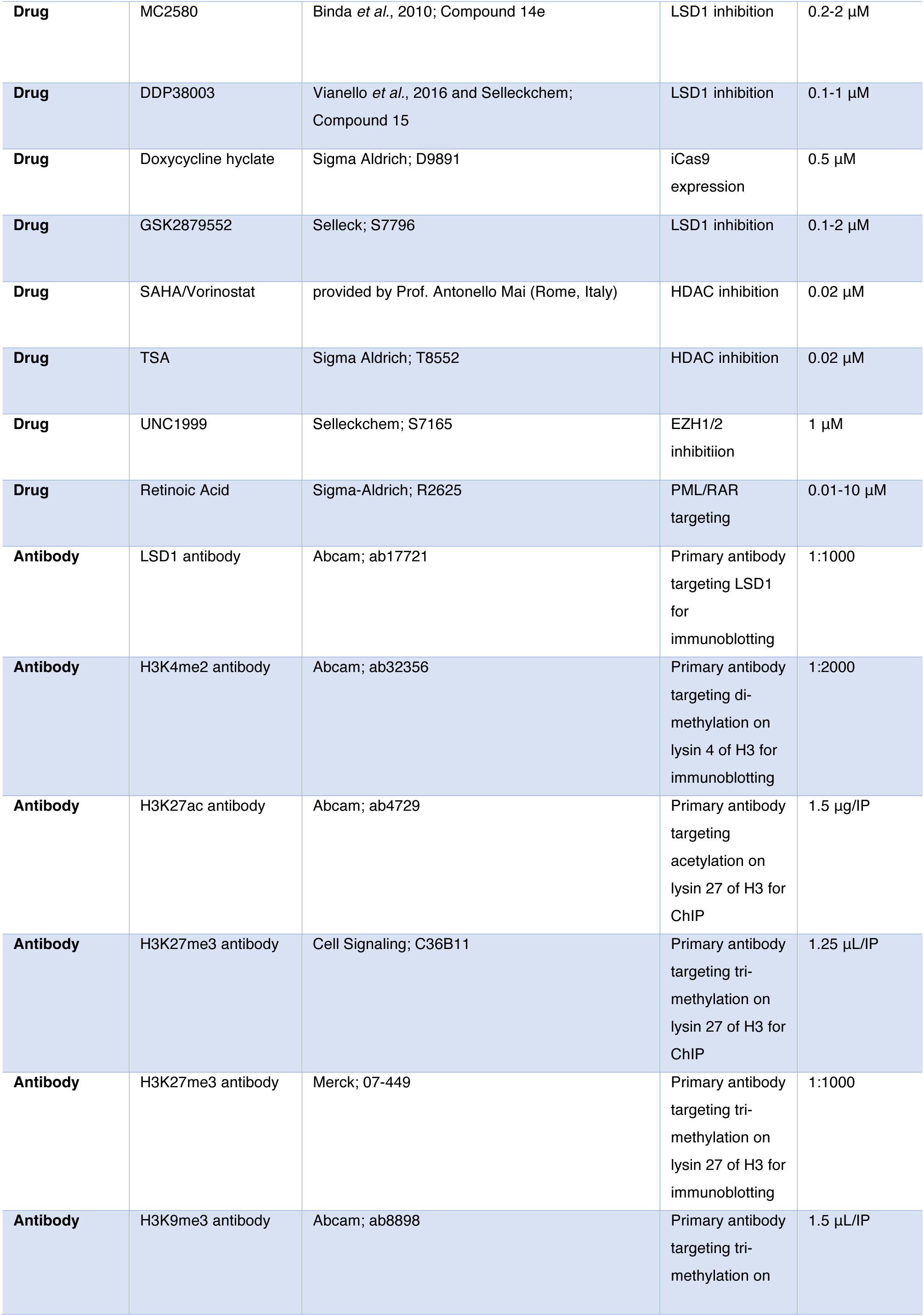
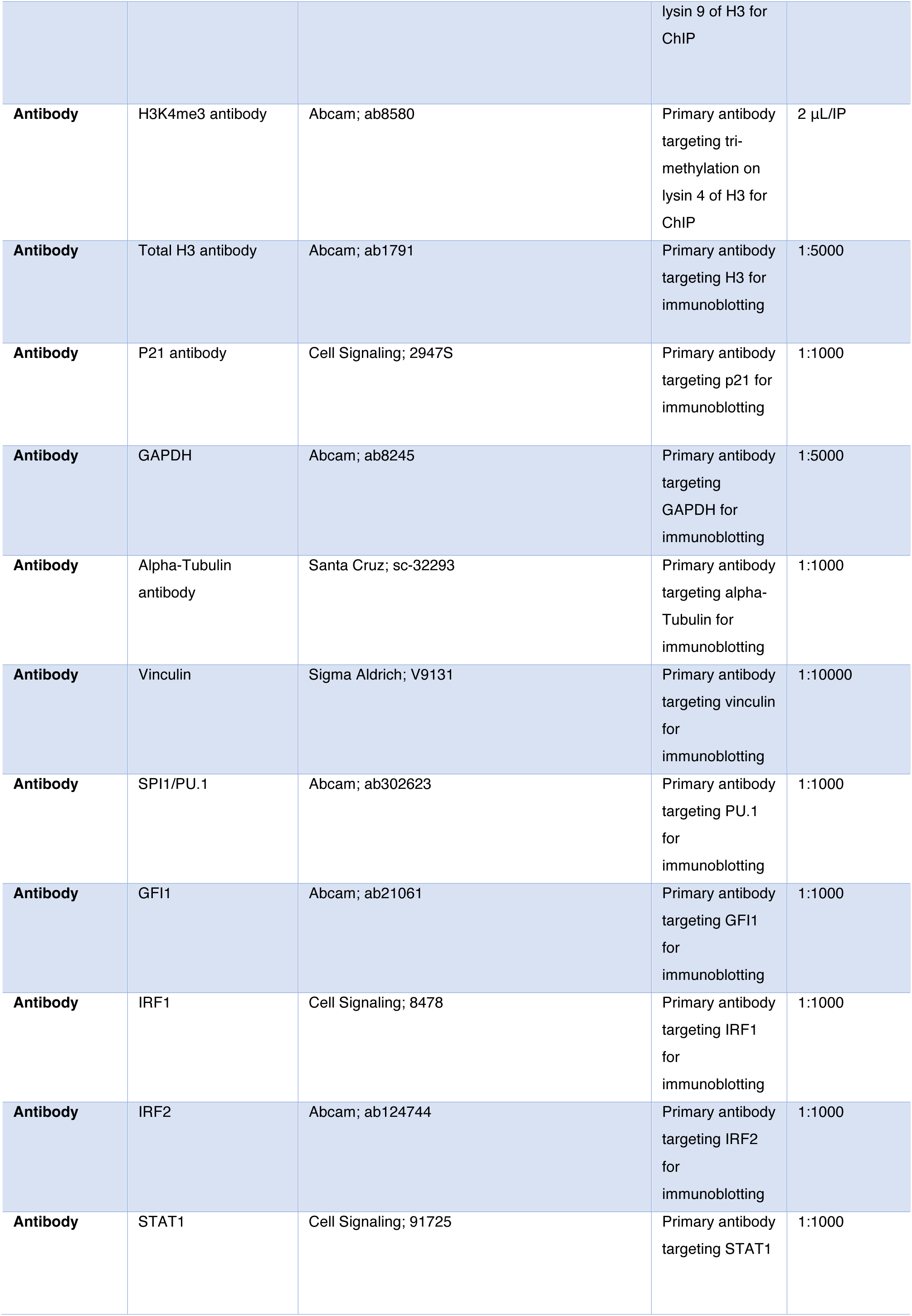
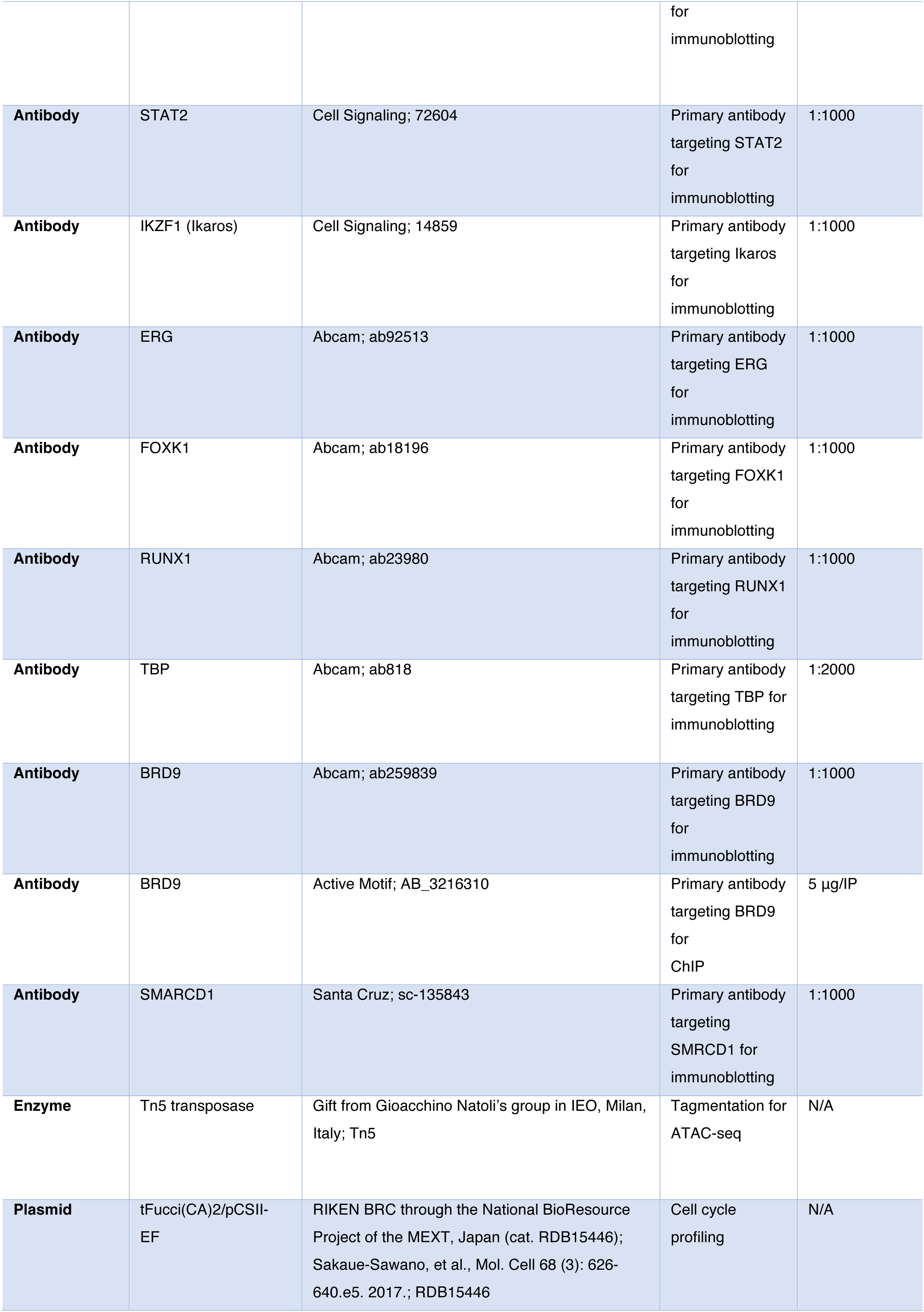
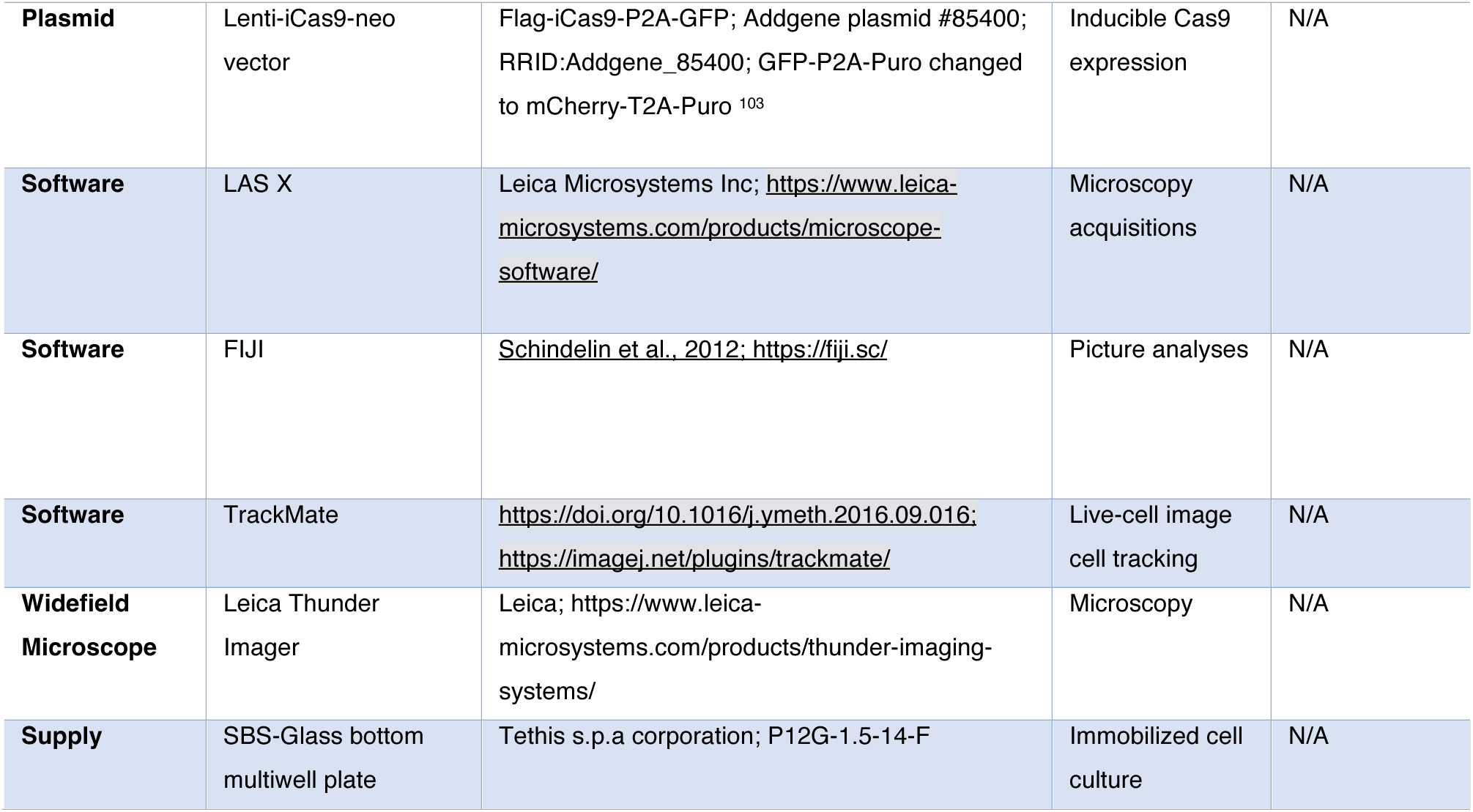
Table of treatments, antibodies, reagents, software and hardware. Details about the compounds used in the screening panel are listed in the supplementary data.

**Table 2.**
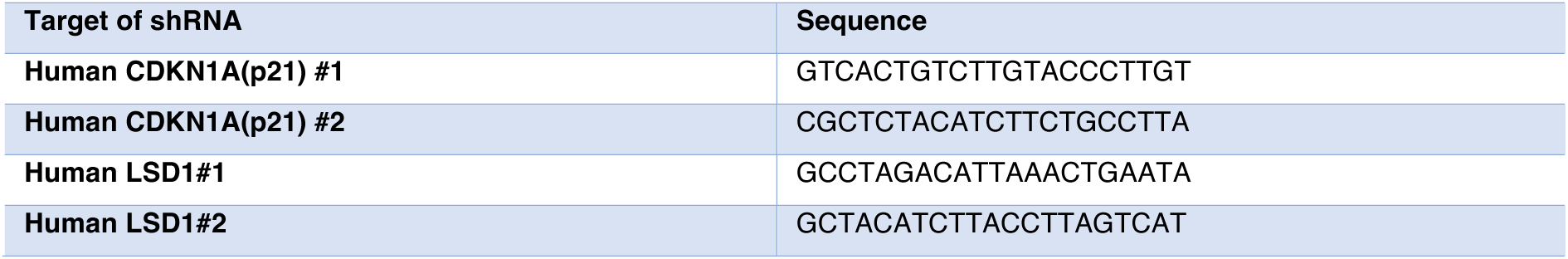
Sequence of oligonucleotide constructs used for RNA interference in this study.

**Table 3.**
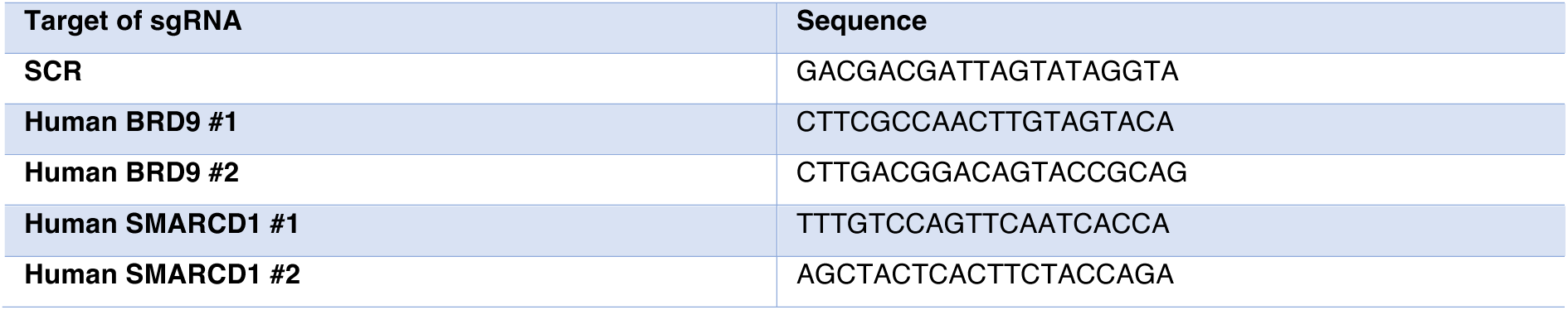
Sequence of oligonucleotide constructs used for sgRNA-mediated KO in this study.

**Table 4.**
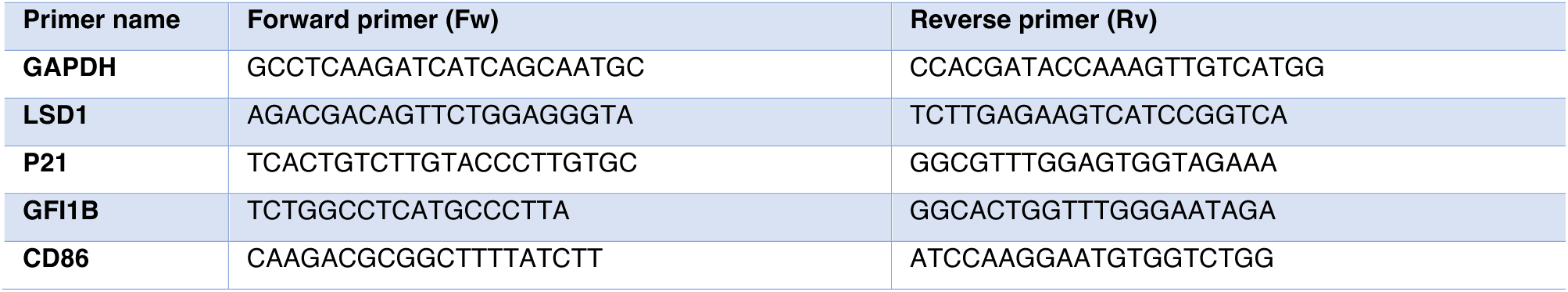
Sequence of primers used for qPCR in this study.

**Table 5.**
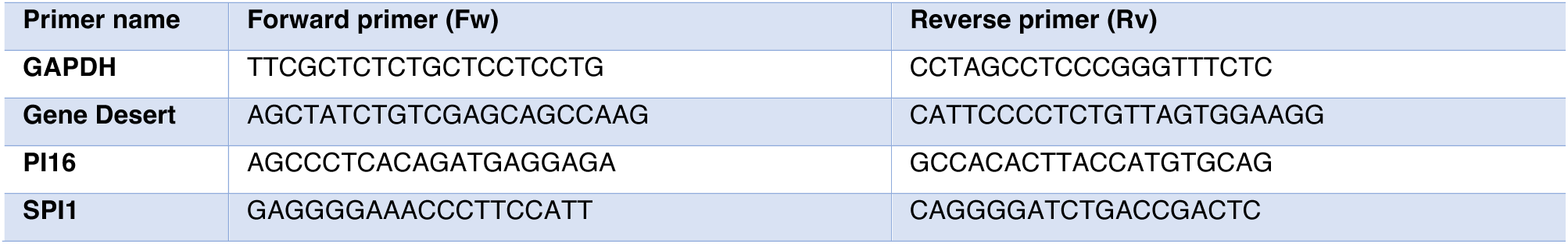
Sequence of primers used for ChIP-qPCR in this study.

## Acknowledgements

The authors are thankful to the scientific and technical support received from Gioacchino Natoli, Claudio Vernieri, Jukka Westermarck, Geneviève Almouzni, Gianmaria Frigè, Fabio Santoro, Marta Kovatcheva, Evelyn Oliva Savoia, and Chiara Ghirardi.

This work was partially supported by a Grant from Fondazione AIRC (IG20 24944) to SM, and partially supported by funds from the Italian Ministry of Health with Ricerca Corrente and 5x1000. KH was supported by Marie Skłodowska-Curie Innovative Training Network (grant no. 813327 ‘ChromDesign’) and AIRC fellowship (ID: 28467).

---

**Extended Data Figure 1.**
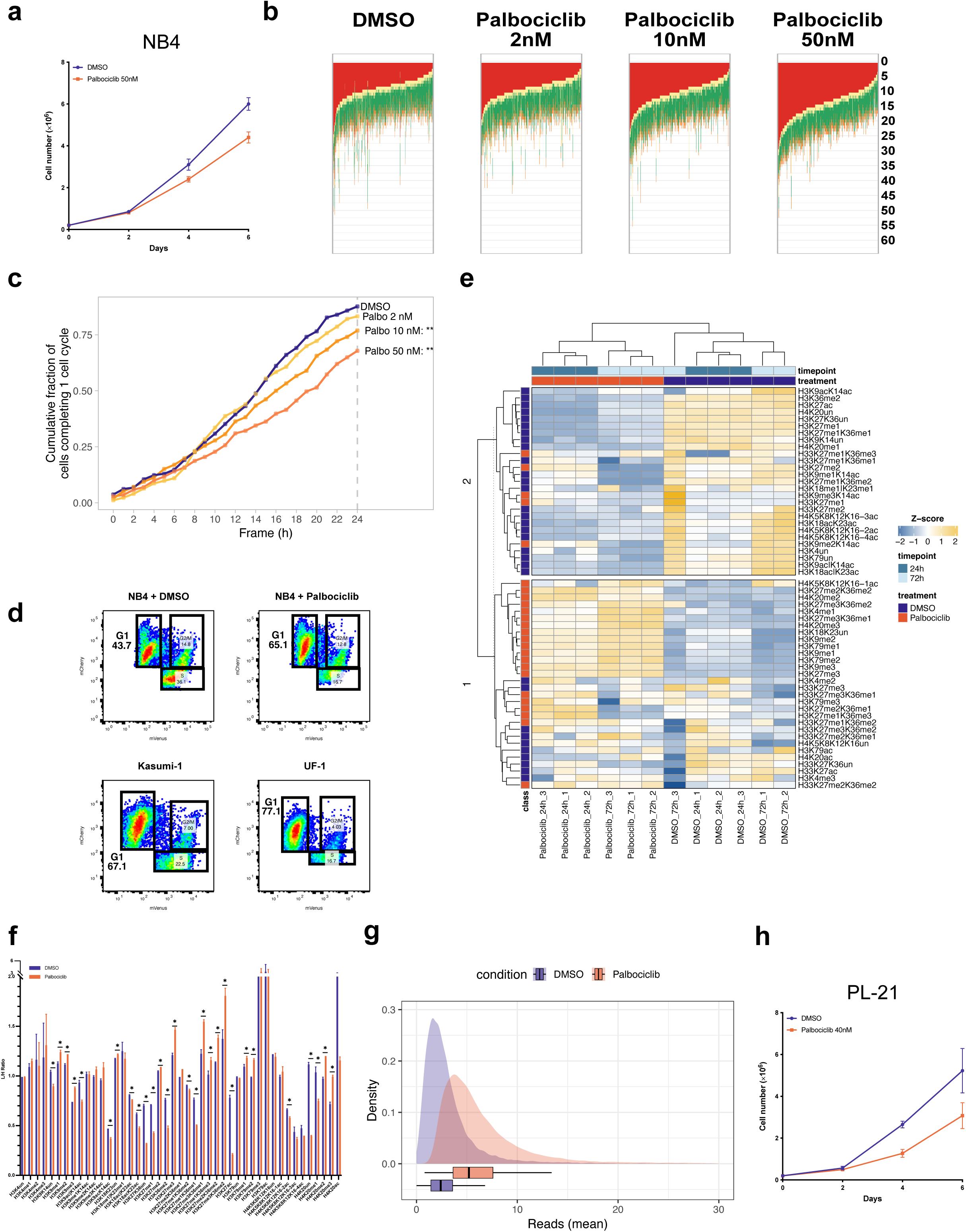
**a**, Growth curve of NB4 cells treated with DMSO or 50 nM Palbociclib. **b**, Waterfall graph representing single cells’ cell cycle phase quantification by live-cell imaging analysis of FUCCI(CA)2-NB4 cells treated with DMSO or Palbociclib at increasing nanomolar concentrations, tracked ours. G1: Red; G1-S transition: Yellow; S: Green; G2/M: Orange. **c**, Line plots showing the cumulative fraction of cells completing one full cycle at different time frames up to 24 hours, normalized to the total number of initial cells. Differences between curves reflect changes in cell cycle progression dynamics induced by the various treatments. Asterisks indicate significant differences relative to the DMSO control (*** P < 0.001). **d**, FACS analysis showing the cell cycle profile of NB4 treated for 24 h with DMSO or 50 nM palbociclib vs. UF-1 and Kasumi-1 based on FUCCI(CA)2 probes expression. **e**, Heatmap of MS-quantified hPTMs in NB4 cells treated with DMSO or Palbociclib (50 nM, 24 and 72 h). Values are mean of log₂[light:heavy] (L/H) ratios from three biological replicates. Rows (hPTMs) and columns (replicates) were hierarchically clustered; rows further partitioned by k-means (k = 2). Annotation bar highlights hPTMs enriched with Palbociclib (orange) or DMSO (dark blue). Heatmap colors denote log₂[L/H] magnitude (blue→yellow). **f**, Expanded bar chart of altered hPTMs (Statistical significance was assessed using an unpaired two-tailed t-test with Welch’s correction. Multiple comparisons were controlled using the two-stage Benjamini–Krieger–Yekutieli FDR procedure, Q = 1%) in NB4 cells as shown in Fig.1c. Bars represent mean L/H ratios (n = 3); error bars indicate replicate variability. Colors: DMSO (blue), Palbociclib (orange). **g**, Density plot showing ATAC-seq reads enrichment in unique Palbociclib peaks shown in Fig.1f. **h**, Growth curve of PL-21 cells treated with DMSO or 40 nM Palbociclib.

**Extended Data Figure 2.**
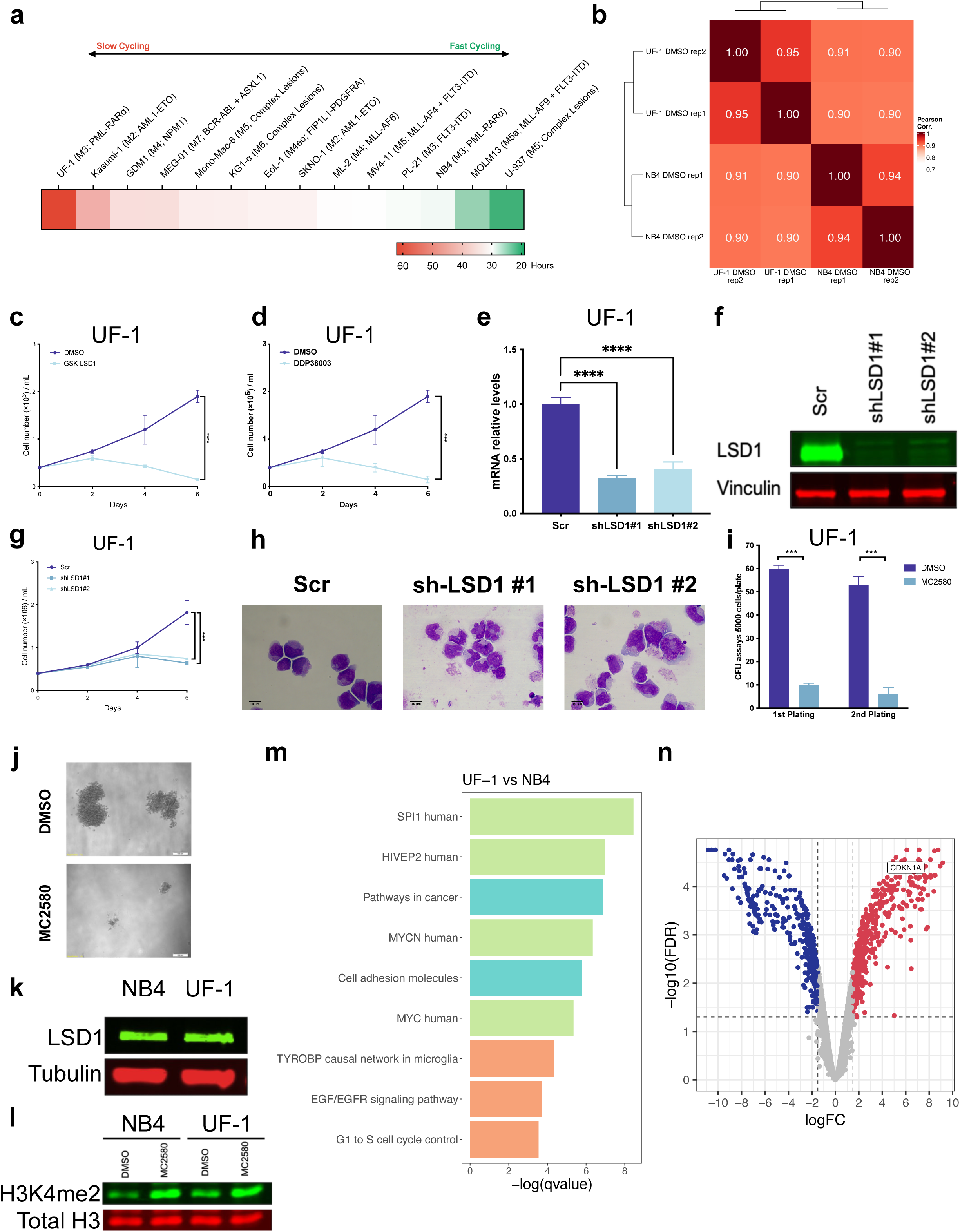
**a,** Heatmap showing doubling times of the panel of AML cell lines shown in ‘Fig.2d’ in normal growth conditions over 6 days of culture. **b,** Heatmap showing transcriptomic correlation between NB4 and UF-1 cells based on RNA-seq study in two biological replicates. **c,** Growth curve (6 days) of UF-1 cells treated with DMSO or GSK-LSD1 (500 nM). Data are mean ± SD of triplicates. **d**, Growth curves (6 days) of UF-1 cells treated with DMSO or DDP38003 (1 µM). Data are mean ± SD of triplicates. **e-f,** Bar chart (e) showing analysis of LSD1 mRNA relative levels and immunoblot (f) showing analysis of LSD1 in UF-1 cells transduced with the indicated lentiviral vectors. Values were normalized against *GAPDH* and referred to Scr. Data are presented as mean ± SD. Statistical significance was assessed by ordinary one-way ANOVA followed by Dunnett’s multiple-comparisons test, with Scr as the control. ns, not significant; ✱P < 0.05, ✱✱P < 0.01, ✱✱✱P < 0.001, ✱✱✱✱P < 0.0001. Vinculin was used as a loading control. **g-h,** Growth curve (g) (6 days) and May-Grünwald-Giemsa-stained images (h) (day 7) of UF-1 cells transduced with the indicated lentiviral vectors. Data are mean ± SD of triplicates. **i-j,** Bar charts (i) showing analysis of the proliferative potential of UF-1 cells treated with DMSO or MC2580 (2 µM). Data are presented as mean of triplicates ± SD. Statistical significance was assessed by ordinary two-way ANOVA followed by [multiple-comparisons test]. ns, not significant; ✱P < 0.05, ✱✱P < 0.01, ✱✱✱P < 0.001, ✱✱✱✱P < 0.0001. Images (j) showing colony morphology of treated UF-1 cells treated with DMSO or MC2580 (2 µM). **k-l,** Western blot analysis of LSD1 (k) and H3K4me2 (l) levels in UF-1 and NB4 cells. Tubulin and total H3 were used as loading controls. **m**, Gene ontology enrichment of DEGs between UF-1 and NB4 cells (FDR < 0.01; |FC| > 1.5). Bars show adjusted P values for enriched terms from KEGG (**Light Blue**), TRRUST (**Light Green**), and WikiPathways datasets (**Light Orange**). **n**, Volcano plot of DEGs between UF-1 and NB4 cells (FDR < 0.01, |FC| > 1.5). Downregulated genes in blue; upregulated in red. All quantitative growth data across the figure were analyzed by one-way ANOVA with Bonferroni’s post-hoc test. P < 0.05 (✱), P < 0.01 (✱ ✱) and P < 0.001 (✱ ✱ ✱).

**Extended Data Figure 3.**
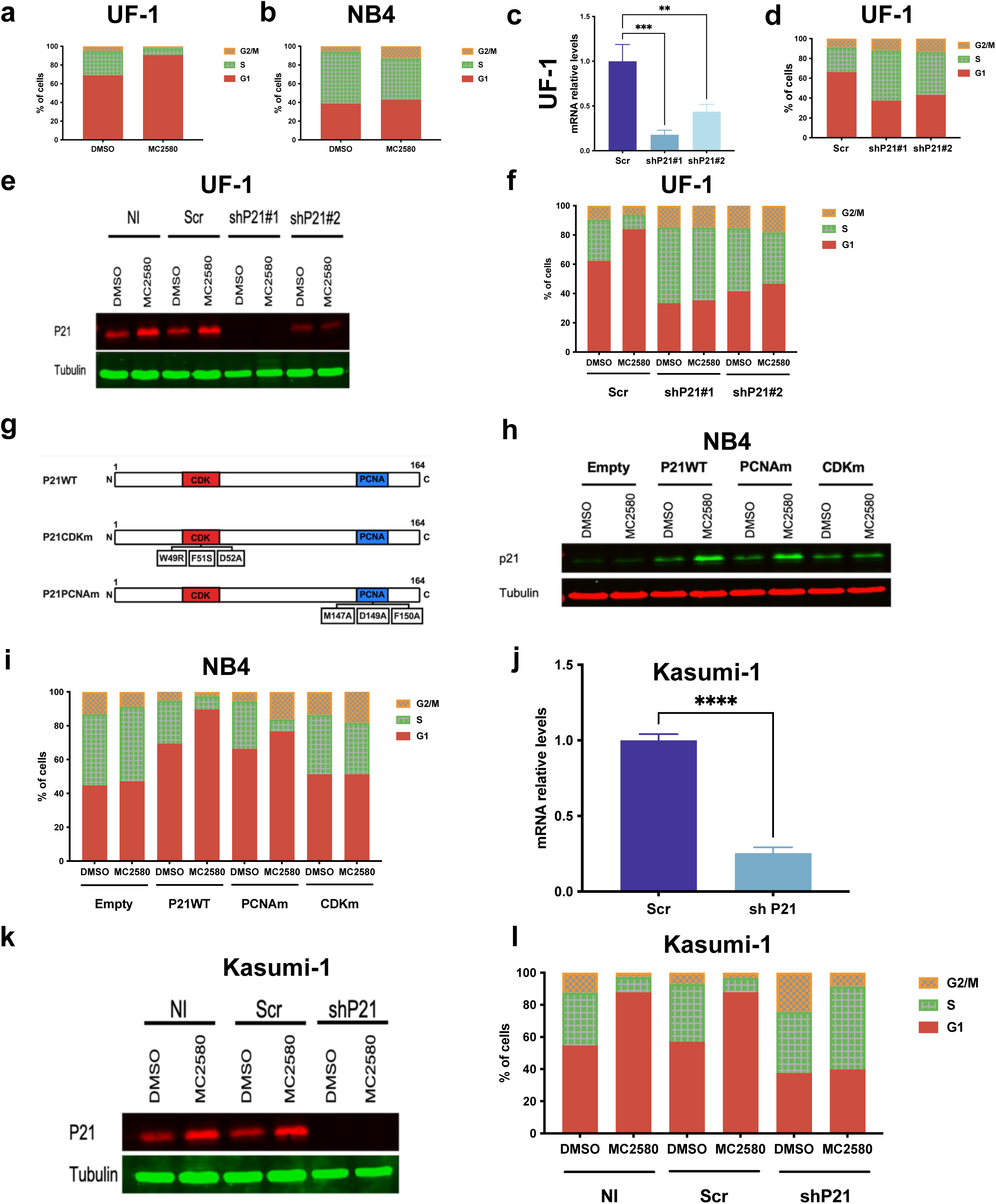
**a-b,** Cell cycle status of UF-1 cells (a) and NB4 cells (b) treated with DMSO or MC2580 (2 µM) for 6 days. **c,** Bar chart showing analysis of *CDKN1A* mRNA relative levels in UF-1 cells transduced with the indicated lentiviral vectors. Values were normalized against *GAPDH* and referred to Scr. Data are presented as mean ± SD. Statistical significance was assessed by ordinary one-way ANOVA followed by Tukey’s multiple-comparisons test. ns, not significant; ✱P < 0.05, ✱✱P < 0.01, ✱✱✱P < 0.001, ✱✱✱✱P < 0.0001. **d**, Cell cycle status of UF-1 cells transduced with either control shRNA (Scr) or shRNA targeting p21. **e**, Western blot analysis of p21 protein in cell lysates from UF-1 cells transduced with the indicated lentiviral vectors or left not infected (NI), and following treatment with DMSO or MC2580 (2 µM). Tubulin was used as a loading control. **f,** Cell cycle status of UF-1 cells transduced with either control shRNA (Scr) or shRNA targeting p21, or left not infected (NI), and following treatment with DMSO or MC2580 (2 µM). **g,** Scheme of the p21 expression constructs used in Fig.2g. p21 CDKm, unable to associate with CDKs, is impaired in its ability to arrest cell cycle, while p21-PCNAm is defective in binding to PCNA. **h,** Western blot analysis of p21 protein in cell lysates from NB4 cells infected with indicated viral vector and treated with DMSO or MC2580 (2 µM). Tubulin was used as a loading control. **i,** Cell cycle status of NB4 cells stably transduced with the indicated vectors and treated with DMSO or MC2580 (2 µM). **j,** Bar chart showing analysis of *CDKN1A* mRNA relative levels in Kasumi-1 cells transduced with the indicated lentiviral vectors. Values were normalized against *GAPDH* and referred to Scr. Data are presented as mean ± SD. Statistical significance was assessed using a two-tailed unpaired t-test with Welch’s correction. ns, not significant; ✱P < 0.05, ✱✱P < 0.01, ✱✱✱P < 0.001, ✱✱✱✱P < 0.0001. **k**, Western blot analysis of p21 protein in cell lysates from Kasumi-1 cells transduced with the indicated lentiviral vectors or left not infected (NI), and following treatment with DMSO or MC2580 (2 µM). Tubulin was used as a loading control. **l,** Cell cycle status of Kasumi-1 cells transduced with either control shRNA (Scr) or shRNA targeting p21, or left not infected (NI), and following treatment with DMSO or MC2580 (2 µM).

**Extended Data Figure 4.**
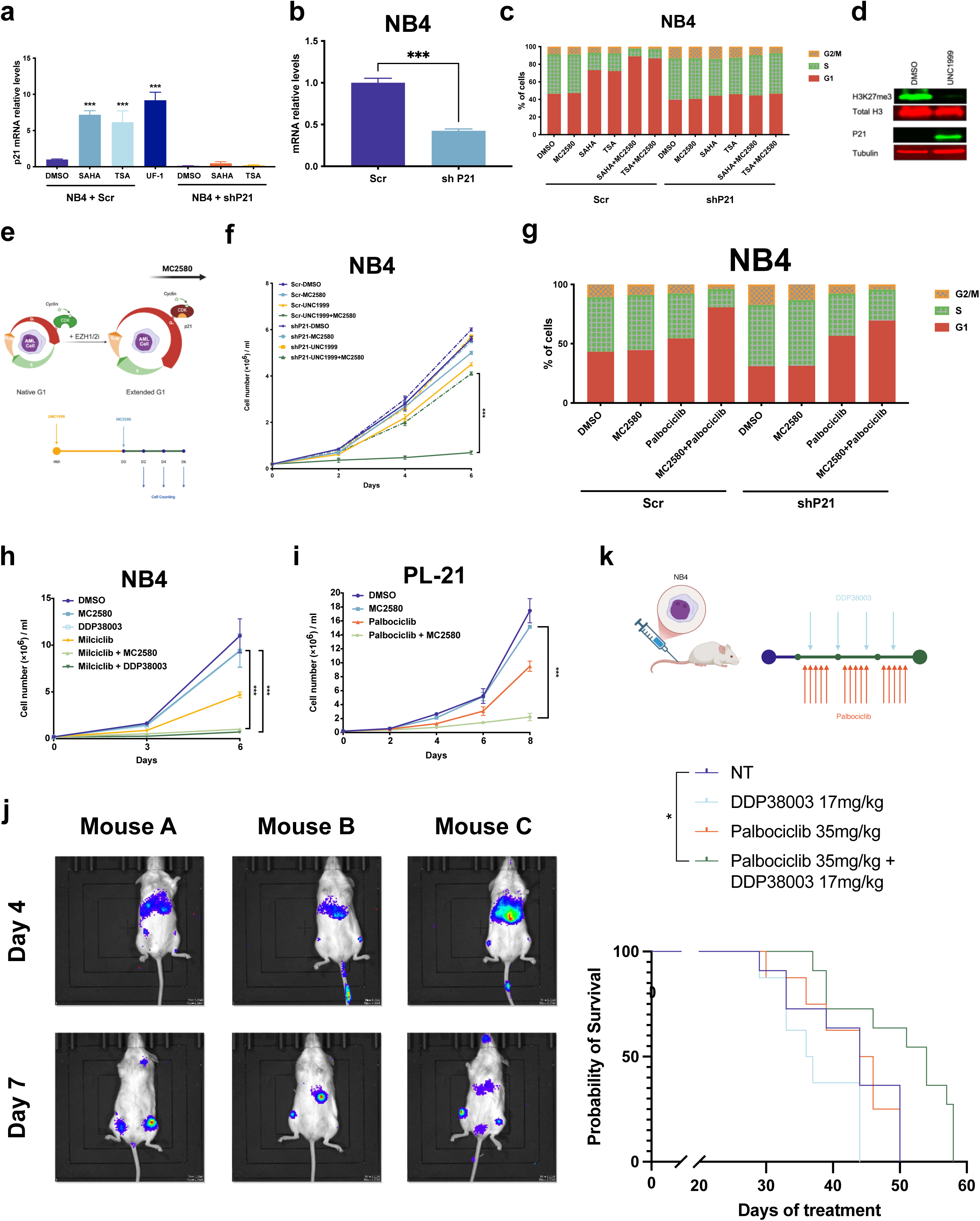
**a,** Bar chart showing analysis of *CDKN1A* (p21) mRNA relative levels in untreated UF-1 cells and NB4 cells transduced with the indicated lentiviral vectors and treated with vehicle (DMSO), HDAC inhibitors (SAHA or TSA, 20 nM). Values were normalized to *GAPDH* and are shown relative to DMSO-treated cells. Relative mRNA levels are presented as mean ± SD. Statistical significance was assessed by ordinary one-way ANOVA followed by Tukey’s multiple-comparisons test. ns, not significant; ✱P < 0.05, ✱✱P < 0.01, ✱✱✱P < 0.001, ✱✱✱✱P < 0.0001. **b,** Bar chart showing *CDKN1A* (p21) mRNA expression in NB4 cells following knockdown with the indicated lentiviral vector and normalized against *GAPDH* referring to Scramble. Data are presented as mean ± SD. Statistical significance was assessed using a two-tailed unpaired t-test with Welch’s correction. ns, not significant; ✱P < 0.05, ✱✱P < 0.01, ✱✱✱P < 0.001, ✱✱✱✱P < 0.0001. **c,** Cell cycle status of NB4 cells stably transduced with the indicated vectors and treated with vehicle (DMSO), MC2580 (2 µM), HDAC inhibitors (SAHA or TSA, 20 nM), or the indicated drug combinations. Data are presented as mean ± SD of triplicates. **d,** Western blot analysis of H3K27me3 and p21 protein levels in NB4 cell lysates. Tubulin and total H3 were used as loading controls. **e,** Experimental scheme for combining UNC1999 with LSD1 inhibition (created with BioRender.com). **f,** Growth curves of NB4 cells treated with the combination of UNC1999 and MC2580. Data are presented as mean ± SD of triplicates. **g,** Cell cycle status of NB4 cells stably transduced with p21 knockdown or control (Scr) vectors and treated with vehicle (DMSO), MC2580 (2 µM), Palbociclib (50 nM), or the indicated drug combination. **h,** Growth curves (6 days) of NB4 cells treated with DMSO, MC2580 (2 µM), Milciclib (500 nM), or the indicated drug combinations. Data are presented as mean ± SD of triplicates. **i,** Growth curves (6 days) of PL-21 cells treated with DMSO, MC2580 (2 µM), Palbociclib (40 nM), or the indicated drug combinations. Data are presented as mean ± SD of triplicates. All quantitative growth data across the previous panels were analyzed by one-way ANOVA with Bonferroni’s post-hoc test. P < 0.05 (✱), P < 0.01 (✱ ✱) and P < 0.001 (✱ ✱ ✱). **j,** Representative whole-animal bioluminescence images of NSG-SGM3 mice injected with 100,000 luciferase-expressing NB4 cells at day 4 and day 7 post-injection. **k,** Top: schematic representation of the *in vivo* experimental design using 100,000 NB4 cells/mouse (experiment workflow created with BioRender.com). Three days post-injection, mice were assigned to the following groups: untreated (NT, 8M+3F), DDP38003 (5M+3F), palbociclib (5M+3F), or DDP38003 + palbociclib (8M+3F). Treatments were administered for 3 weeks, and mice were monitored and sacrificed upon onset of clinical signs. Bottom: Survival of the indicated mice over the course of treatment. Survival differed significantly among groups (Mantel–Cox log-rank test, P = 0.0010). Pairwise comparisons against NT were performed with correction for multiple comparisons using the Holm–Šídák method. Adjusted P values: NT vs DDP38003, P = 0.1594; NT vs palbociclib, P = 0.8476; NT vs combination, P = 0.0275. *P < 0.05; ns, not significant. NT, n = 11; DDP38003, n = 8; palbociclib, n = 8; DDP38003 + palbociclib, n = 11 mice.

**Extended Data Figure 5.**
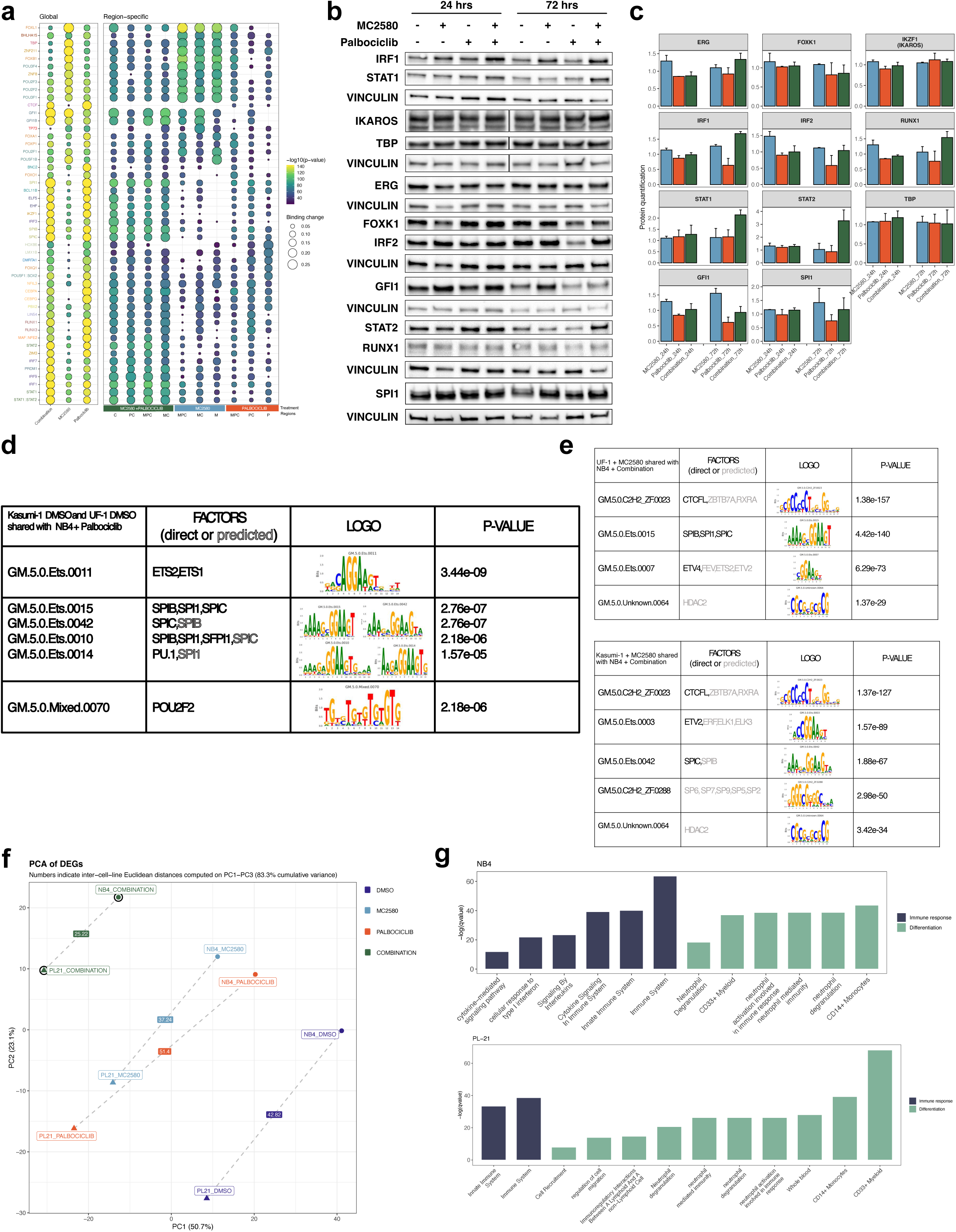
**a,** Bubble plot showing TOBIAS footprinting comparing TF binding in NB4 cells treated with Palbociclib + MC2580, Palbociclib, or MC2580 versus DMSO (day 1). The top 10 TFs for each treatment and region-specific comparison were selected using the most significant motif per TF. Regions referring to Fig. 4a are indicated by the same initials. Circle size indicates differential binding and color intensity reflects −log10(P-value); negative differential binding values are not shown. **b-c,** Western blot analysis (b) of selected transcription factors in NB4 cells treated with MC2580 (2 µM), Palbociclib (50 nM), their combination or vehicle (DMSO) for 24 and 72 h, with corresponding quantification of protein levels normalized to Vinculin and DMSO (c) in at least two biological replicates. **d,** Table showing TOBIAS motif discovery of top TF binding sites identified in mutually open chromatin regions based on ATAC-seq analysis in G1-sorted untreated Kasumi-1, UF-1 and 50 nM palbociclib-treated NB4 cells. **e,** Table showing TOBIAS motif discovery of top TF binding sites identified in mutually open chromatin regions based on ATAC-seq analysis in G1-sorted 2 µM MC2580-treated Kasumi-1, UF-1 and 50 nM palbociclib + 2 µM MC2580-treated NB4 cells. **f,** PCA showing transcriptomic correlation of NB4 and PL-21 cells treated for 24 h with DMSO, MC2580 (2 µM), Palbociclib (50 and 40 nM, respectively), or their combination. The analysis was based on DEGs discovered in at least one condition. **g,** Bar plots showing gene ontology enrichment of DEGs in NB4 (up, MC2580 + Palbociclib vs DMSO) and PL-21 (down, MC2580 + Palbociclib vs DMSO, day 1). Bars show adjusted P values for top terms (Reactome, Human Gene Atlas, GO biological process).

**Extended Data Figure 6.**
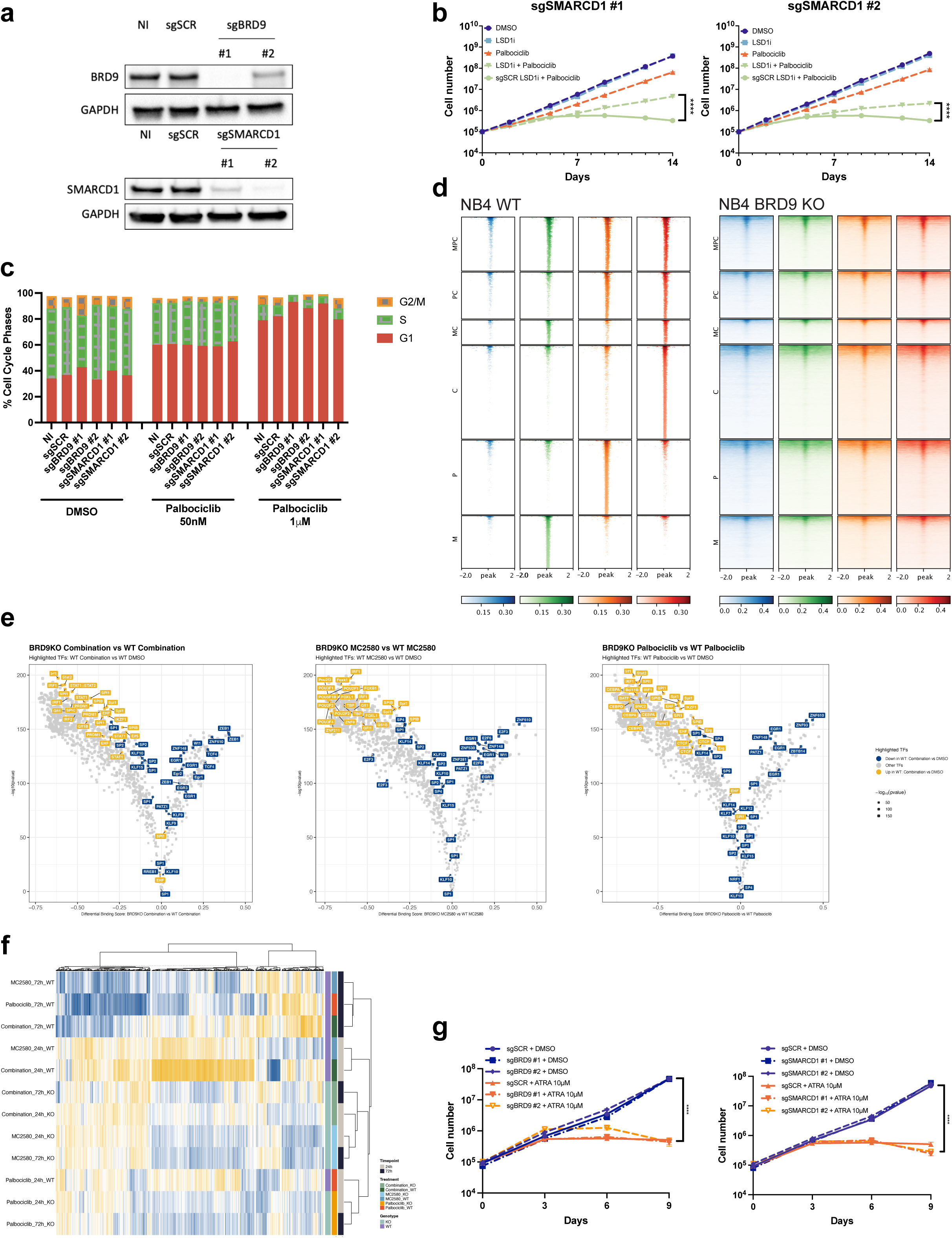
**a,** Western blot analysis of BRD9 (left) and SMARCD1 (right) proteins in cell lysates from NB4 cells transduced with the indicated lentiviral vectors or left not infected (NI). **b**, Growth curves (14 days) of sgRNA-expressing NB4 cells treated with DMSO, MC2580 (2 µM), Palbociclib (50 nM), or combination shown in Log_10_ scale. Data are mean ± SD of triplicates. **c**, FACS analysis using DAPI staining showing the cell cycle profile of NB4 cells transduced with the indicated lentiviral vectors or left not infected (NI) treated for 24 h with DMSO, MC2580 (2 µM), Palbociclib (50 nM), or combination. **d,** Tornado plots of regions shown in Figure 4a in WT (left) and BRD9-KO (right) NB4 cells treated with DMSO, MC2580 (2 µM), Palbociclib (50 nM), or their combination. **e,** Volcano plots showing differential TF footprinting between BRD9-KO and WT NB4 cells under the indicated treatments (2 μM MC2580, 50 nM palbociclib, or their combination for 24 h). Each point represents a transcription factor motif, plotted by differential binding score and –log10 P value. TFs previously identified among the top 20 positively (yellow) or negatively (blue) differential footprints in the corresponding treatment vs DMSO comparison in WT cells are highlighted and annotated **f,** Heatmap of RNA expression for the top 811 significantly upregulated genes shown in Figure 4d in WT (normal line) and BRD9-KO (dotted line) NB4 cells treated with palbociclib (50 nM), MC2580 (2 µM), or their combination versus DMSO (day 3). **g,** Growth curves (9 days) of sgRNA-expressing NB4 cells treated with DMSO, or ATRA (10 µM) in Log_10_ scale. Data are mean ± SD of triplicates. All quantitative growth data across the figure were analyzed by two-way ANOVA with Tukey’s multiple comparisons test. P < 0.05 (✱), P < 0.01 (✱ ✱) and P < 0.001 (✱ ✱ ✱). Analysis of the data in panel b was performed on log_10_-transformed values.

**Extended Data Figure 7.**
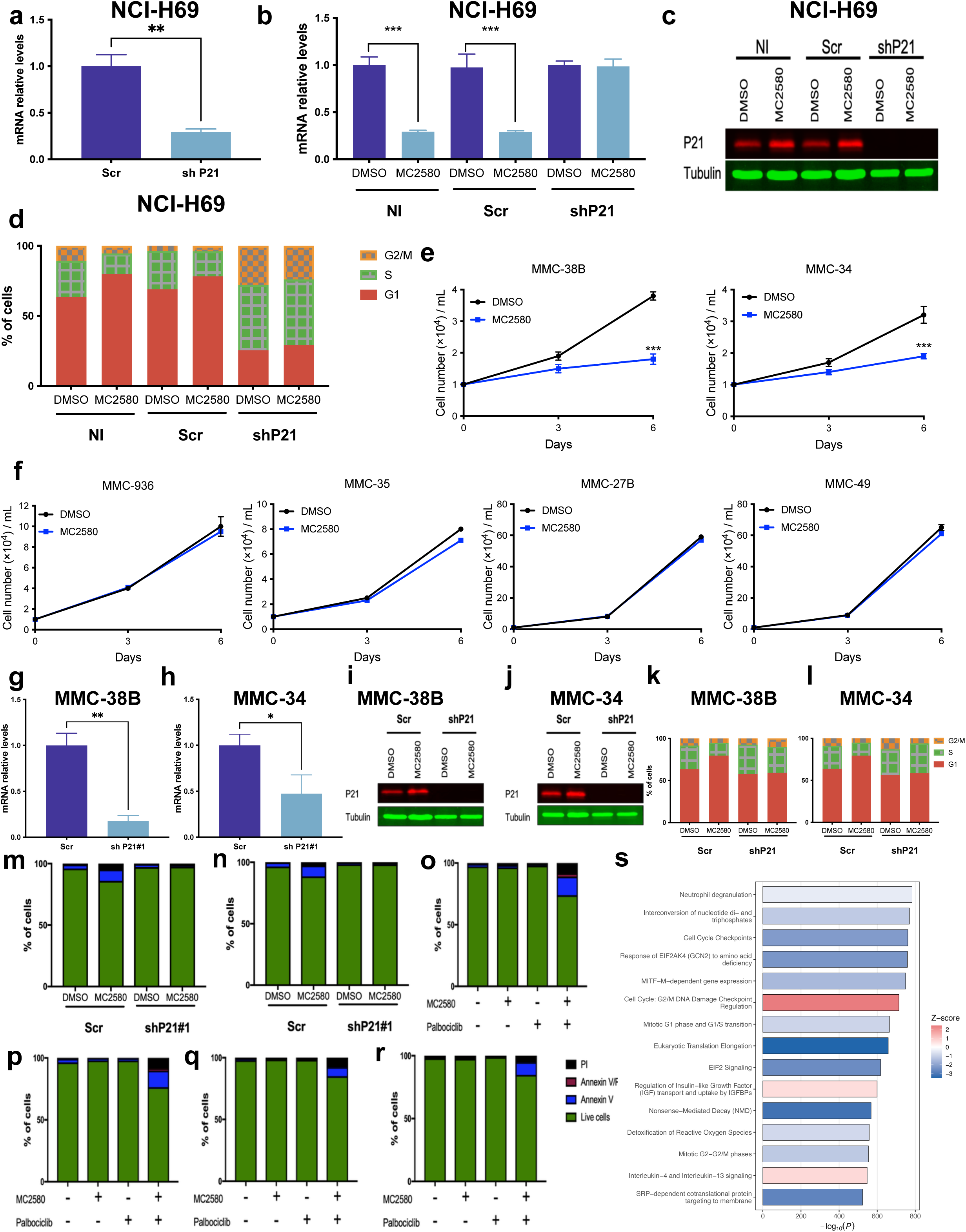
**a,** Bar chart showing *CDKN1A* (p21) mRNA relative levels in NCI-H69 cells transduced with the indicated lentiviral vectors. Values were normalized to *GAPDH* and are shown relative to Scr control. Data are presented as mean ± SD. Statistical significance was assessed using a two-tailed unpaired t-test with Welch’s correction. ns, not significant; ✱P < 0.05, ✱✱P < 0.01, ✱✱✱P < 0.001, ✱✱✱✱P < 0.0001. **b,** Bar chart showing *GRP* mRNA (immature cell marker) relative levels in NCI-H69 cells stably transduced with the indicated vectors or left non-infected (NI), following treatment with vehicle (DMSO) or MC2580 (2 µM) for 6 days. Data are presented as mean ± SD. Statistical significance was assessed by ordinary one-way ANOVA followed by Tukey’s multiple-comparisons test. ns, not significant; ✱P < 0.05, ✱✱P < 0.01, ✱✱✱P < 0.001, ✱✱✱✱P < 0.0001. **c,** Western blot analysis of p21 protein in NCI-H69 cell lysates as in Figure 6f. Tubulin was used as a loading control. Cell-cycle distribution of NCI-H69 cells as in Figure 6f. **d,** Cell cycle status of NCI-H69 cells stably transduced with p21 knockdown or control (Scr) vectors or left not-infected (NI) and treated with vehicle (DMSO) or MC2580 (2 µM). **e-f,** Growth curves of LSD1i-sensitive (e) and LSD1i-insensitive (f) human primary melanoma cells treated with vehicle (DMSO) or MC2580 (2 µM) for 6 days. Data are presented as mean ± SD of triplicates. **g-h,** Bar charts showing *CDKN1A* (p21) mRNA relative levels in MMC-38B (g) and MMC-34 (h) cells transduced with the indicated lentiviral vectors. Values were normalized to *GAPDH* and are shown relative to Scr control. Data are presented as mean ± SD. Statistical significance was assessed using a two-tailed unpaired t-test with Welch’s correction. ns, not significant; ✱P < 0.05, ✱✱P < 0.01, ✱✱✱P < 0.001, ✱✱✱✱P < 0.0001. **i,** Western blot analysis of p21 protein in MMC-38B cell lysates as in Figure 6h (left). Total H3 was used as a loading control. **j,** Western blot analysis of p21 protein in MMC-34 cell lysates as in Figure 6h (right). Total H3 was used as a loading control. **k-l,** Cell cycle status of MMC-38B (k) and MMC-34 (l) cells transduced with the indicated lentiviral vectors. **m-n,** Percentage of live and apoptotic MMC-38B (m) and MMC-34 (n) cells transduced with the indicated lentiviral vectors and treated with vehicle (DMSO) or MC2580 (2 µM). Legends are shown in panel “r.” **o-r,** Percentage of live and apoptotic MMC-936 (o), MMC-35 (p), MMC-49 (q) and MMC-27B (r) cells treated with vehicle (DMSO), MC2580 (2 µM), Palbociclib (50 nM), or their combination. Legends are shown in panel “r.” **s**, IPA of DEGs in WM266.4 DTPs cells versus parental cells.

